# Experimental alteration of the microRNA profile of the mouse embryo: heritable phenotypes in the adult

**DOI:** 10.1101/2023.02.21.529218

**Authors:** Zeynep Yilmaz Sukranli, Kezban Korkmaz Bayram, Serpil Taheri, Francois Cuzin, Yusuf Ozkul, Minoo Rassoulzadegan

**Affiliations:** Betul-Ziya Eren Genome and Stem Cell Center, Erciyes University, Kayseri, Turkey; Department of Medical Genetics, Faculty of Medicine, Erciyes University, Kayseri, Turkey; Department of Medical Genetics, Faculty of Medicine, Yıldırım Beyazıt University, Ankara, Turkey; Department of Medical Biology, Faculty of Medicine, Erciyes University, Kayseri, Turkey; INSERM-CNRS, Universite de Nice, Nice, France

## Abstract

We previously identified a unique genetic feature of Autism Spectrum Disorder (ASD) in human patients and established mouse models, namely the low to very low level of six microRNAs, miR-19a-3p, miR-361-5p, miR-3613-3p, miR-150-5p, miR-126-3p and miR-499a-5p. We attempted to interfere experimentally with the expression in the mouse of two of them, miR19a-3p and miR499a-5p by microinjecting into the one-cell embryo either the complementary sequence or an excess of the microRNA. Both approaches resulted in low levels in the somatic tissues and sperm of the targeted microRNAs (nascent) and their pri and pre precursors. The altered miRNA profiles were inherited by progenies of crosses with untreated partners together with behavior alterations partly characteristic of ASD patients and animal models. An excess of miRNA in the eggs leads to a specific downregulation, we propose variations of single-stranded miRNA, as effectors of its own transcription in eukaryotes.

## Introduction

Searching for the genetic determinant(s) of Autism Spectrum Disorder (ASD) uncovered a large number of variations in patient genomes, variable associations of partial determinants of the whole complex phenotype including de novo mutations^1–4^. Most of these genetic alterations were, however found in only a fraction of the patients and thus could not be primary determinants of the disease. The first and only exception to date has been our observation of a deviant pattern of microRNAs in the blood of 45 autistic patients^5^ in a cohort of 37 families. The levels of six of them, miR-19a-3p, miR-361-5p, miR-3613-3p, miR-150-5p, miR-126-3p and miR-499a-5p, were more than 95 percent down-regulated in serum levels of affected patients children and were reduced by about 50 % in their healthy parents and siblings. Further studies have extended the analysis to two established animal models of ASD, mice treated with valproic acid and *Cc2d1a+/-* heterozygotes. In both cases, the strongly decreased levels of miR-19a-3p, miR-361-5p, miR-150-5p, miR-126-3p and miR-499a-5p initially observed in the blood of human patients, were evidenced in the blood, hippocampus and sperm of affected animals. In addition, the animal model allowed us to establish that altered miRNA profiles were inherited together with ASD-like symptoms in crosses with unaffected animals.

We hypothesized that the altered miRNA pattern of the progenitors might act as a transgenerational determinant. While miRNAs have mainly been studied as regulators of mRNA translation^6^, several of them were reported to act as epigenetic regulators, positively or negatively interfering, with the transcription of target genes^7^. We attempted to deregulate miRNAs profiles by microinjection of homologous oligonucleotides into one-cell mouse embryos. For a first of the tests, we considered miR-19a-3p as a candidate based on an in silico search of its possible targets which indicated no less than five mouse loci whose human counterpart has been associated with ASD, *Pten, Slc6a4*, *Fmr1*, *Foxp2* and *Cc2d1a*^1, 8–12^. miR-499a-5p, singled out by the family analysis^5^, was also retained. The availability of a faithful animal model of the disease has enabled us to respond positively to two critical questions, namely whether experimentally in mice we can modify in a stable way the levels of predetermined miRNAs and in the case of ASD, whether it may result in a hereditary disease.

## Results

### Changing the amount of a given miRNA from the egg to the mouse: miRNAs microinjection into fertilized mouse eggs results in complex but reproducible heritable changes in adult tissues

To check whether microinjection of a microRNA into a fertilized egg would result in a stably altered expression in the adult, we first examined animals derived from eggs microinjected with the complementary sequence of the two microRNAs under study (Figure 1 and Supplementary Table 1). We had initially hypothesized that microinjection of an excess of the complementary sequence or of the miRNA itself would result in their either decrease or increase in the copy number. Since the resulting patterns turned out to be reproducibly found in animals derived from eggs independently microinjected and their offspring, this unexpected result actually provide us in fact with a useful tool for a more in-depth analysis.

**Figure 1:**
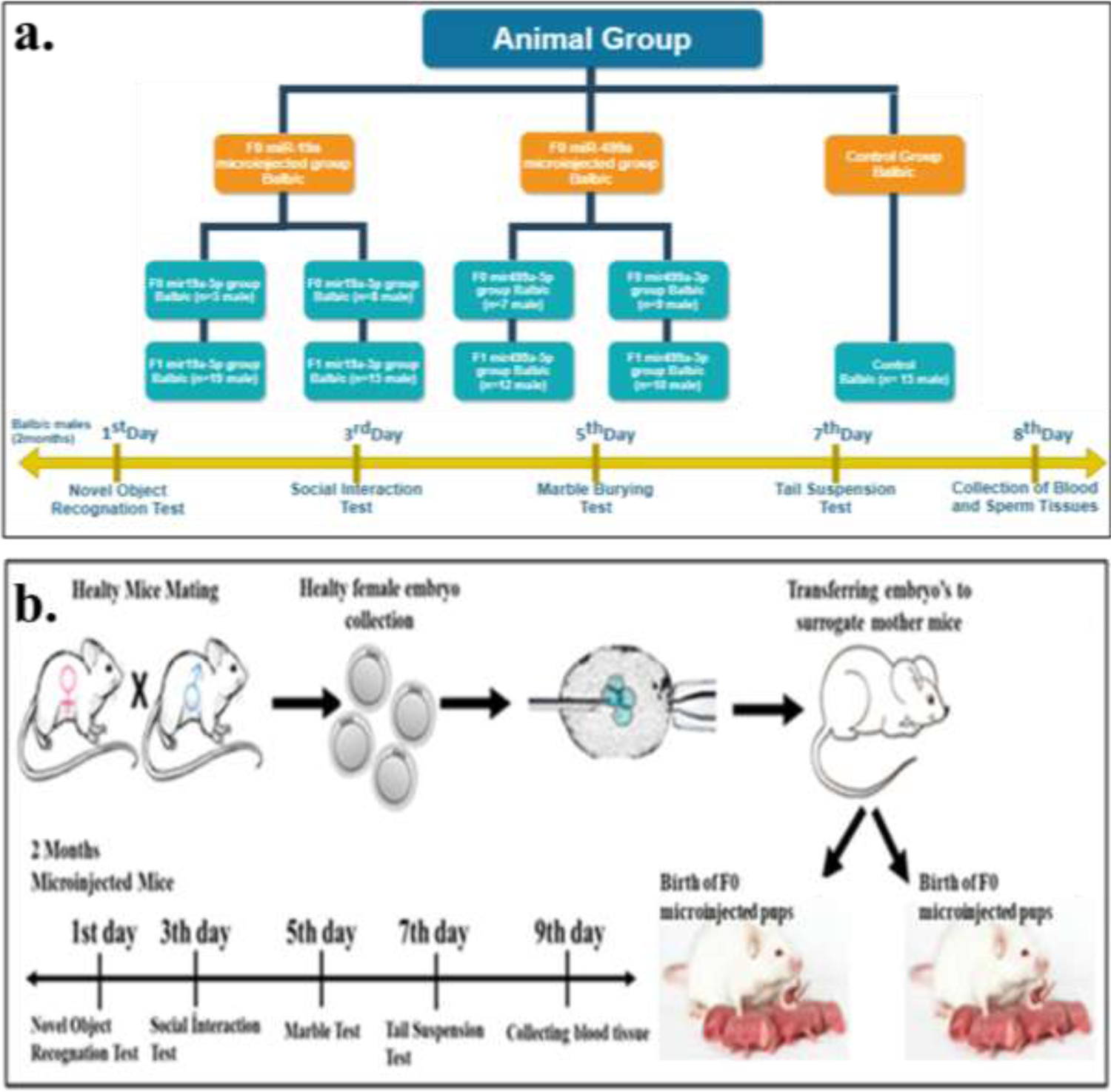
**a.** Molecular and behavioral tasks timeline **b.** Demonstration of microinjection procedure

A summary of mice born after microinjection and miRNAs analysis is listed in Figure 1.

The expression levels of six miRNAs after microinjection of a specific miRNA are presented in Figure 2 and 3. The first series of observations evidenced the expected decrease after microinjection of the complementary sequence but also by that of an excess of miRNA.

**Figure 2:**
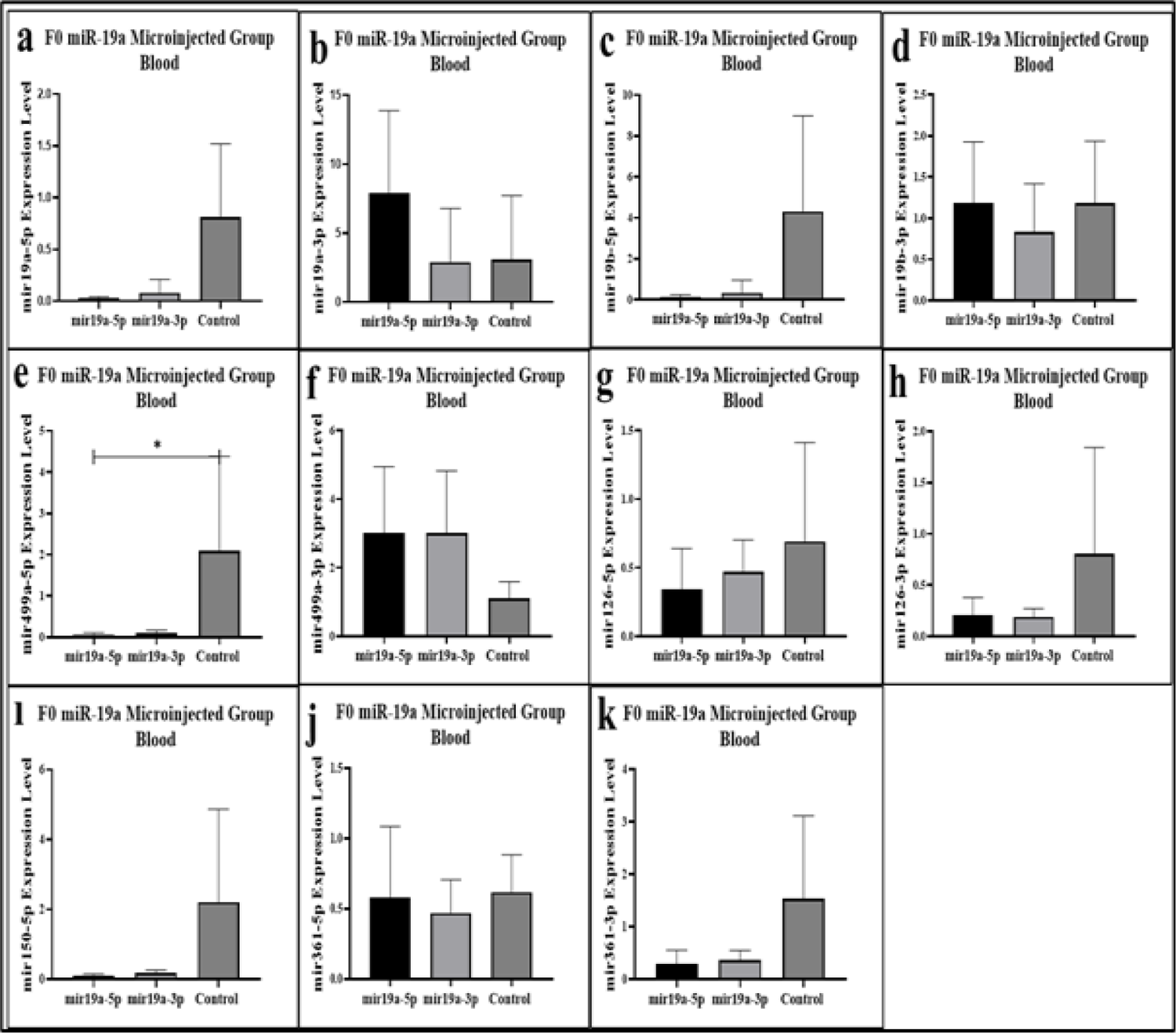
miRNAs expression levels results in the blood of miR-19a microinjected mice. **a.** miR-19a-5p expression levels, **b.** miR-19a-3p expression levels, **c.** miR-19b-5p expression levels, **d.** miR-19b-p expression levels, **e.** miR-499a-5p expression levels, **f.** miR-499a-3p expression levels, **g.** miR-126a-5p expression levels, **h.** miR-126a-3p expression levels, **ı.** miR-150a-5p expression levels, **j.** miR-361a-5p expression levels and **k.** miR-361a-3p expression levels.

**Figure 3:**
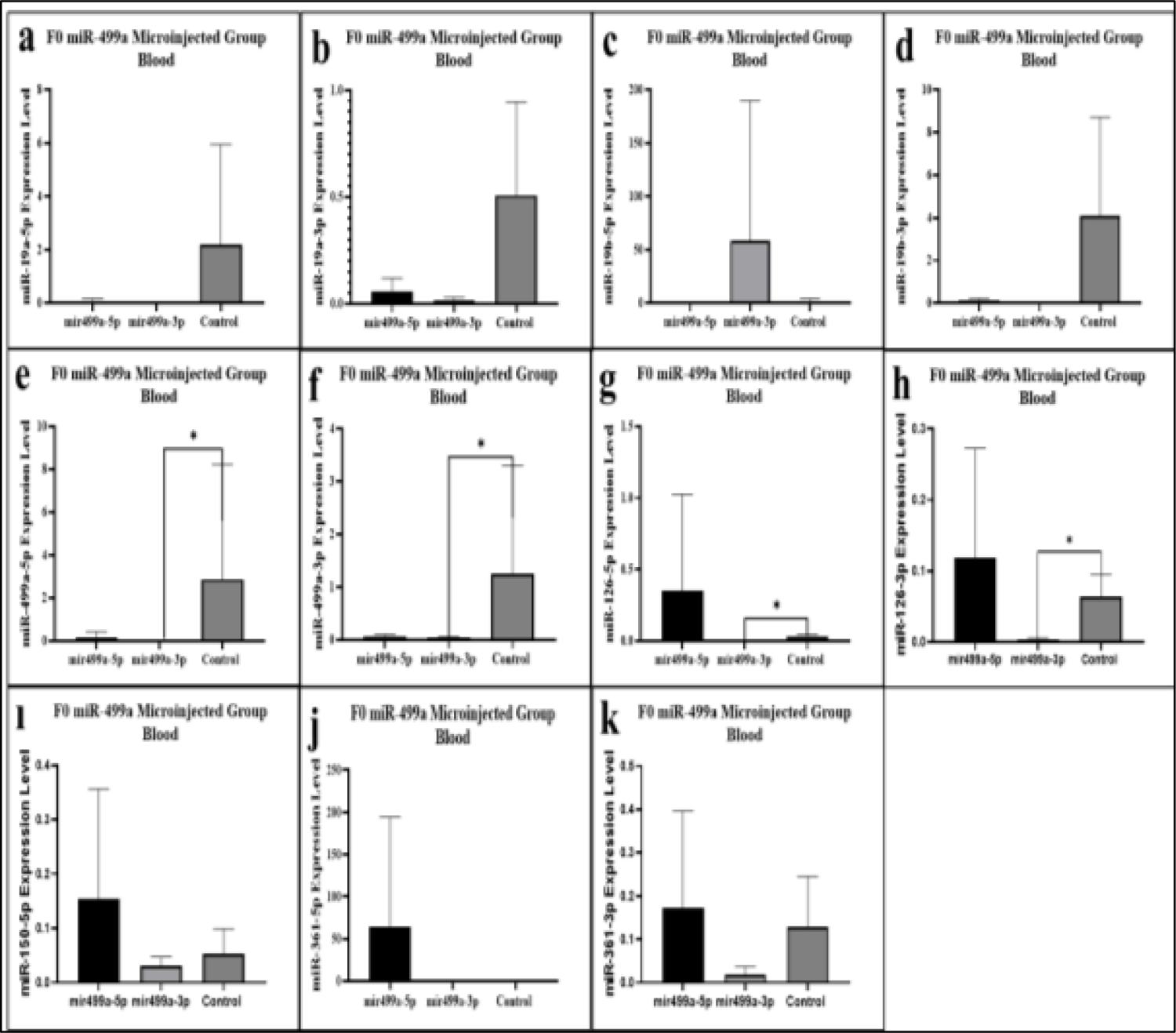
miRNAs expression levels results in the blood of miR-499a microinjected mice (5p/3p). **a.** miR-19a-5p expression levels, **b.** miR-19a-3p expression levesl, **c.** miR-19b-5p expression levels, **d.** miR-19b-p expression levels, **e.** miR-499a-5p expression levels, **f.** miR-499a-3p expression levels, **g.** miR-126a-5p expression levels, **h.** miR-126a-3p expression levels, **ı.** miR-150a-5p expression levels, **j.** miR-361a-5p expression levels and **k.** miR-361a-3p expression levels.

Altogether our results indicate that miRNAs microinjection modifies (their) own expression and could influence not only its reverse sequences but also other microRNA levels. For example miR-19a and miR-19b are transcribed from a polycistronic locus^13^. Microinjection of miR-19a and b (−5p and −3p) was found to down-regulate miR-19a-5p transcripts levels, while miR-19a-3p and miR-19b-3p are unchanged by miR-19a-3p and or upregulated by the miR-19a-5p strand (Figure 2). Of the other miRNAs-(see Table 1) miR-499-5p, miR-150-5p, miR-126-3p and miR-361-3p are down-regulated and miR-499-3p is up-regulated (Figure 2).

In mice microinjected with miR-499a-3p, four microRNAs were downregulated compared to controls: miR-499a-3p, miR-499a-5p, miR-126-3p, miR-126-5p. The microinjection results validate the multiple effect of miR19a-3p on miR499-5p (up-regulation) and the subsequent effect of miR-499a-5p and miR-499a-3p on its own transcriptions and mainly on miR-126-3p, miR-126-5p (Figure 3).

These results support our first assumption that exposure of one-cell stage embryos to miRNAs permanently affects miRNAs levels (here five mi*RNA*s) in mouse blood cells.

### Down-regulation at the transcription level

The micro*RNA*s are transcribed by RNA polymerase II (RNAP II), and the products (pri-mi*RNA*s) are converted into pre-mi*RNA*s by Drosha-DGCR8. Next, we examined the level of immature mi*RNA*s transcripts. At this point, our analyses used separate specific oligonucleotides (see Supplementary Table 2 for oligonucleotide sequences) and RT q-PCR to amplify transcript fragments specific for Pri and Pre miRNAs accumulation. As shown in Figure 4 and 5 from our results in blood and Supplementary Figures in the hippocampus (see in Supplementary Figure 1 for mature miRNA expression; see in Supplementary Figure 2 for expression of pri and pre miRNA), miRNAs transcripts Pri and Pre are altered in all tissues after microinjection of mature mi*RNA*s into fertilized eggs. The modification of the levels of the Pri and Pre transcripts supports the notion of a regulatory mechanism perpetuating the signals from one to other, at least for the transcripts disclosed here. Is there a common regulation of transcription, the answer to this important question requires further investigations.

**Figure 4:**
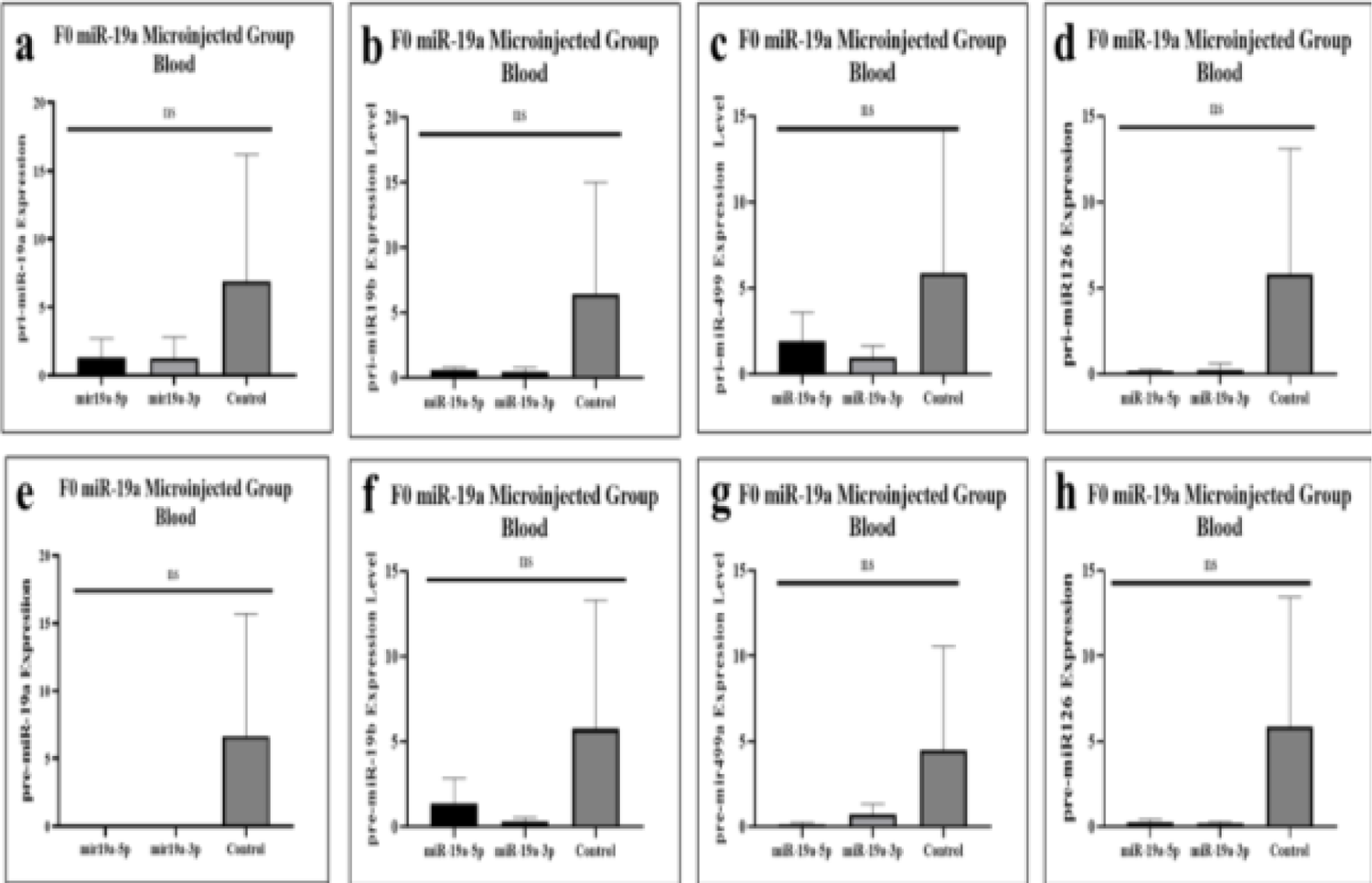
pri and pre-miRNAs’ expression levels results in the blood of miR-19a microinjected mice. **a.** pri-miR-19a expression levels, **b.** pri-miR-19b expression levels, **c.** pri-miR-499a expression levels, **d.** pri-miR-126a expression levels **e.** pre-miR-19a expression levels, **f.** pre-miR-19b expression levels, **g.** pre-miR-499a expression levels and **h.** pre-miR-126a expression levels.

**Figure 5:**
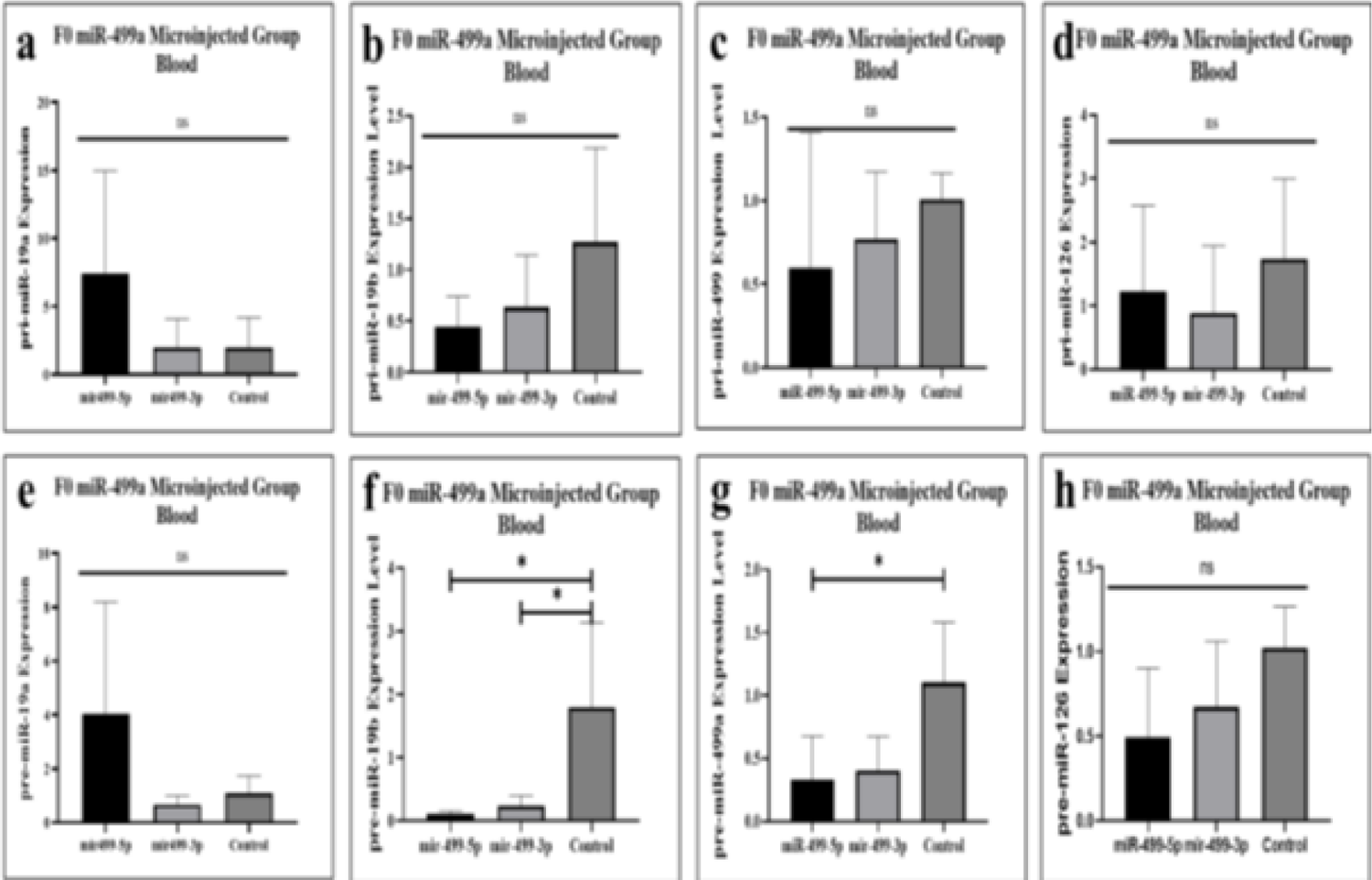
pri and pre-miRNAs’ expression levels results in the blood in miR-499a microinjected mice. **a.** pri-miR-19a expression levels, **b.** pri-miR-19b expression levels, **c.** pri-miR-499a expression levels, **d.** pri-miR-126a expression levels **e.** pre-miR-19a expression levels, **f.** pre-miR-19b expression levels, **g.** pre-miR-499a expression levels and **h.** pre-miR-126a expression levels.

### Does a defect in miRNA distribution play a causal role in behavior? behavior analysis reveals modified animals

We have previously established that, in human patients and animal models, down-regulation of miR-19a-3p, miR-361-5p, miR-150-5p, miR-126-3p and miR-499a-5p is part of component switching behavioral. At that time, it was not possible to establish whether the microRNA defect could in itself be a sufficient causal element in the initiation of behavioral variations. Since we are now able to alter the profile of miRNAs in otherwise healthy organisms, it becomes possible to determine whether altering miRNA one-by-one triggers at least part of the behavioral changes.

Behavior tasks were performed on male mice from 2 months of age through recognition of objects novel to familial, social interactions, tail suspension, marble burying assays (see timeline in Figure 1).

Together all results of the behavioral tasks are shown in Figures 6 and 7, with a number of variations observed with mainly certain less active groups of mice. Thus, the loss of interest between familiar objects and new objects seems to indicate a common deficiency of the “autistic group” of mice comparable to the positive control group of male mice treated with Valproic acid previously shown^5^.

**Figure 6:**
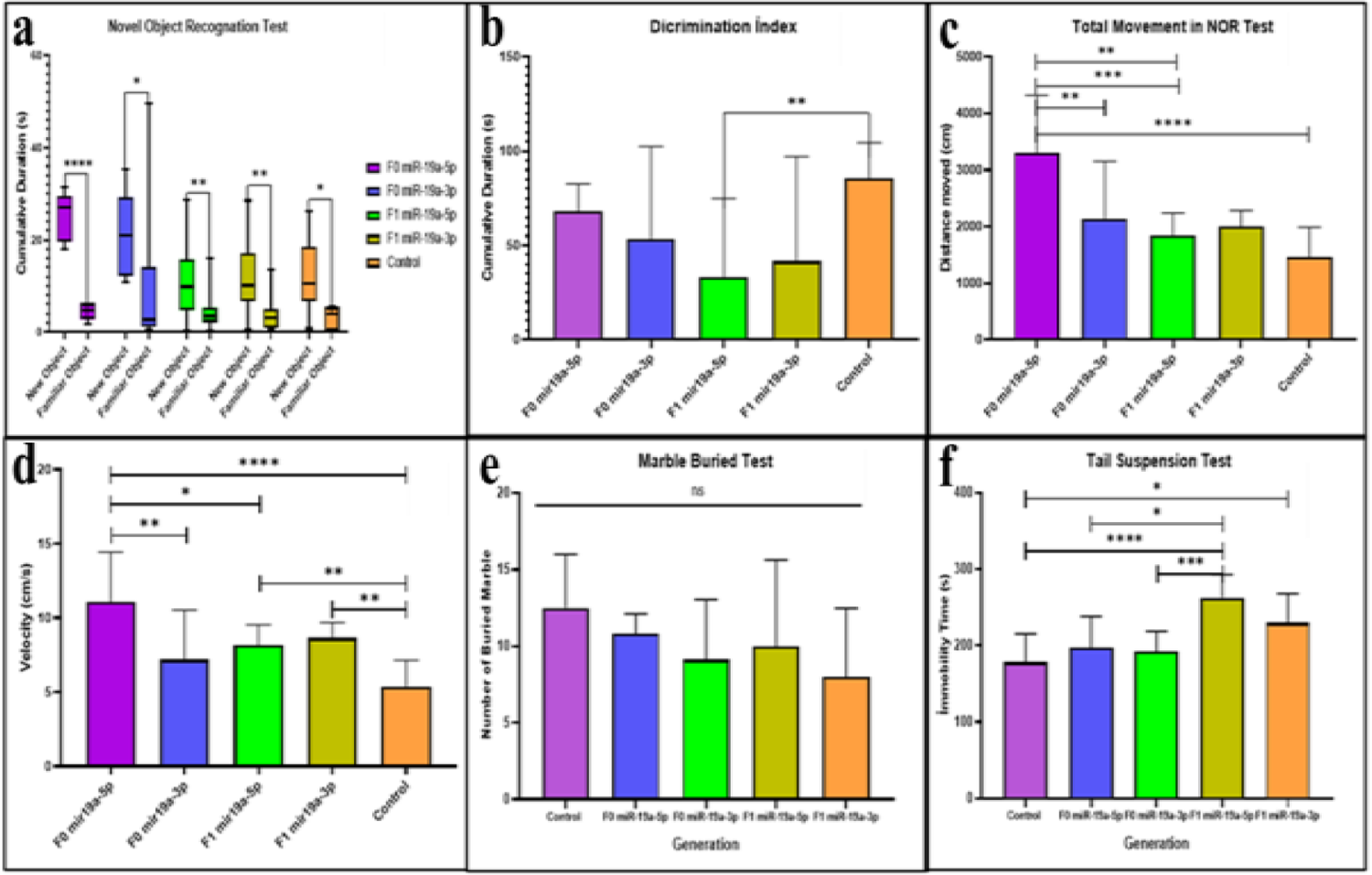
The results of Behaviour tests of miR-19a microinjected group (5p/3p) **a.** Comparison of total exploration time between new object and familiar object, **b.** Comparison of total exploration time in F0 and F1 generations in NOR, **c.** Comparison of total movement between F0 and F1 generations in NOR test, **d.** Comparison of total speed in NOR test, **e.** Comparison of total buried number in F0 and F1 generations in Marble Test and **f.** Comparison of immobility time in F0 and F1 generations in Tail Suspension Test.

**Figure 7:**
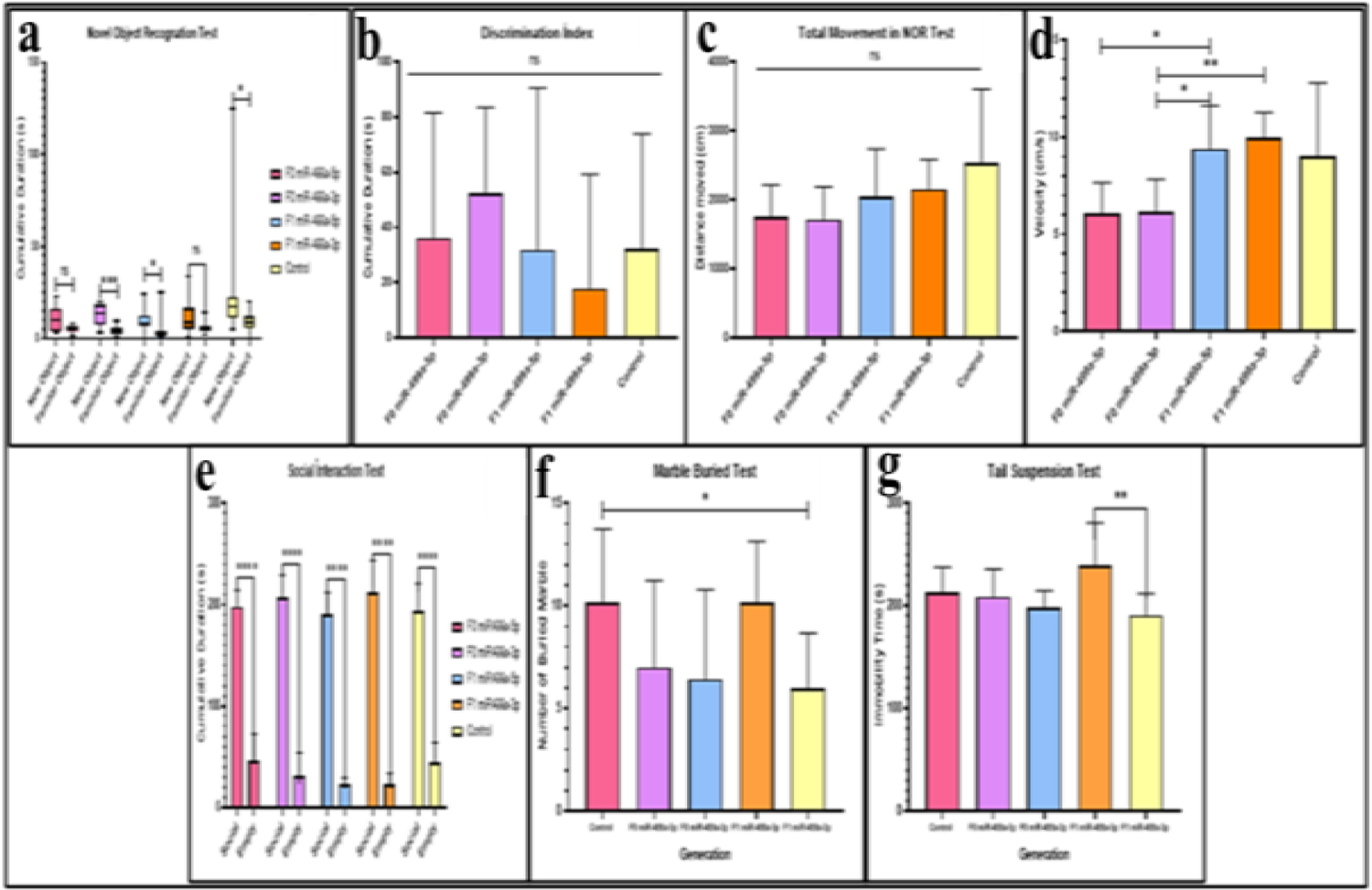
Behaviour tests result of miR-499a microinjected group (5p/3p) **a.** Comparison of total exploration time between the new object and familiar object, **b.** Comparison of total exploration time in F0 and F1 generations in NOR **c.** Comparison of total movement between F0 and F1 generations in NOR test, **d.** Comparison of total speed in NOR test, **e.** Comparison of total social time, **f.** Comparison of total buried number in F0 and F1 generations in Marble Test and **g.** Comparison of immobility time in F0 and F1 generations in Tail Suspension Test.

As shown in Figure 6a interest in the novel object increased in founder mice microinjected with both strands of miR-19a-5p and miR-19a-3p, while their F1 generation showed a comparable phenotype to controls. In contrast, mice born after being microinjected with each strand of miR-499a-5p and miR-499a-3p, now compared to the control group showed reduced interest in both objects observed for the founder as well as the F1 generation (Figure 7a). Mice born after microinjection of miR-499a-5p and miR-499a-3p into fertilized eggs appear to behave with an autism-like phenotype comparable to autistic mouse models as we have previously reported with males treated with Valproic acid^5^.

The discrimination index, which measures the percentage of time spent with the new object, shows that F1 miR-19a-3p spent significantly less time than the control. In addition, we see that F0 miR-19a-5p spent less time than the other groups, while the time spent with the new object was longer than the other groups. Similarly, F0 miR-19a-5p was found to be significantly more mobile and faster than all other groups. Even though there was a decrease in velocity in the F1 miR-19a-5p and miR-19a-3p groups, it was observed that there was a significant increase compared to the control. In the group microinjected with miR-499a, there was no significant difference between the groups in terms of discrimination index and total movement, while the group with the highest discrimination index was F0 miR-499a-3p. This confirms the time spent with the new object. In terms of speed, F1 miR-19a-5p and miR-19a-3p groups were observed to move faster than F0 miR-499a-3p and all other groups.

Self-grooming or balls-burying is frequently observed in the autistic mouse model. However, there were no or only very slight differences in self-grooming and burrowing between the groups of mice microinjected with miR-19a and miR-499a.

The Tail Suspension experiment showed that autistic model mice develop a sedentary posture after the first escape movements when placed in an unavoidably stressful situation. Except mice microinjected with miR-19a, on the other hand, all groups were more motile than control, while only F1 showed significant difference between miR-19a-3p and miR-19a5p. Less mobility is observed in F0 miR-19a-3p within groups. In mice microinjected with miR-499a-3p, F1 shows less activity than miR-499a-5p (see in Figure 7). F1 miR-499a-3p is the least mobile of all groups. There is no significant difference between the other groups. In contrast to the novel object recognition test, the F1 miR-499-3p tail suspension test displayed more autistic behavior.

In conclusion, functional assays validate RNA-mediated alteration of behavioral phenotypes.

### Either a defect in the distribution of miRNA transmitted to the next generation: behavioral analysis reveals modified animals

The expression levels of six miRNAs in the sperm of males born after microinjection of a specific miRNA are shown in Figure 8. The results of sperm miRNAs analysis indicate that microinjection of miRNAs (miR-499a-5p and −3p) dramatically reduced the expression of miR-19a-5p and miR-499a-3p, these were found downregulated in our previous studies in serum from patients with autism and mouse models. Increasing levels of these miRNAs by microinjection in fertilized eggs leads to reduced levels in mouse sperm. This negative regulation is also maintained in the serum of the F1 generation and accompanied by behavioral variations see Figure 6.

**Figure 8:**
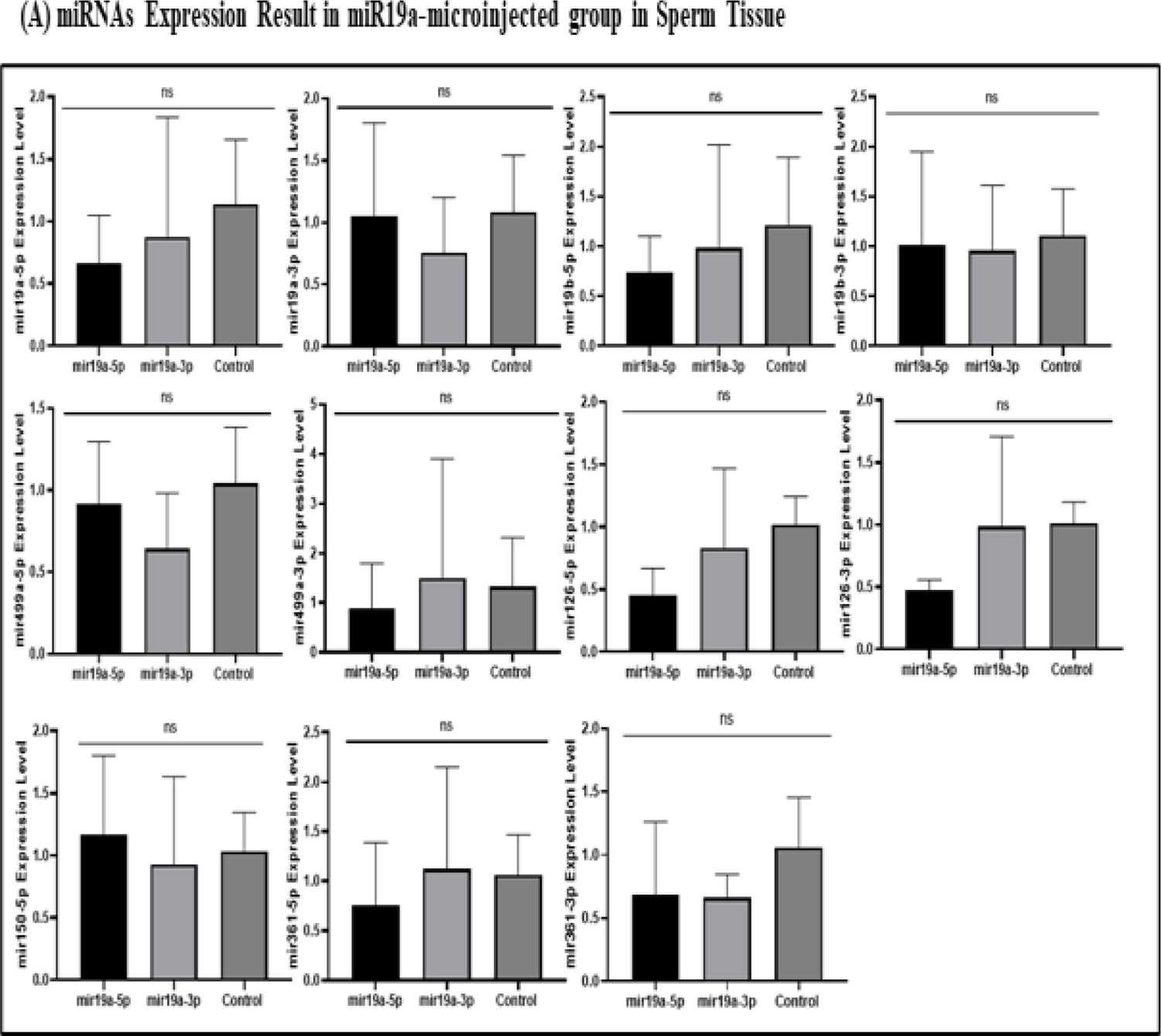
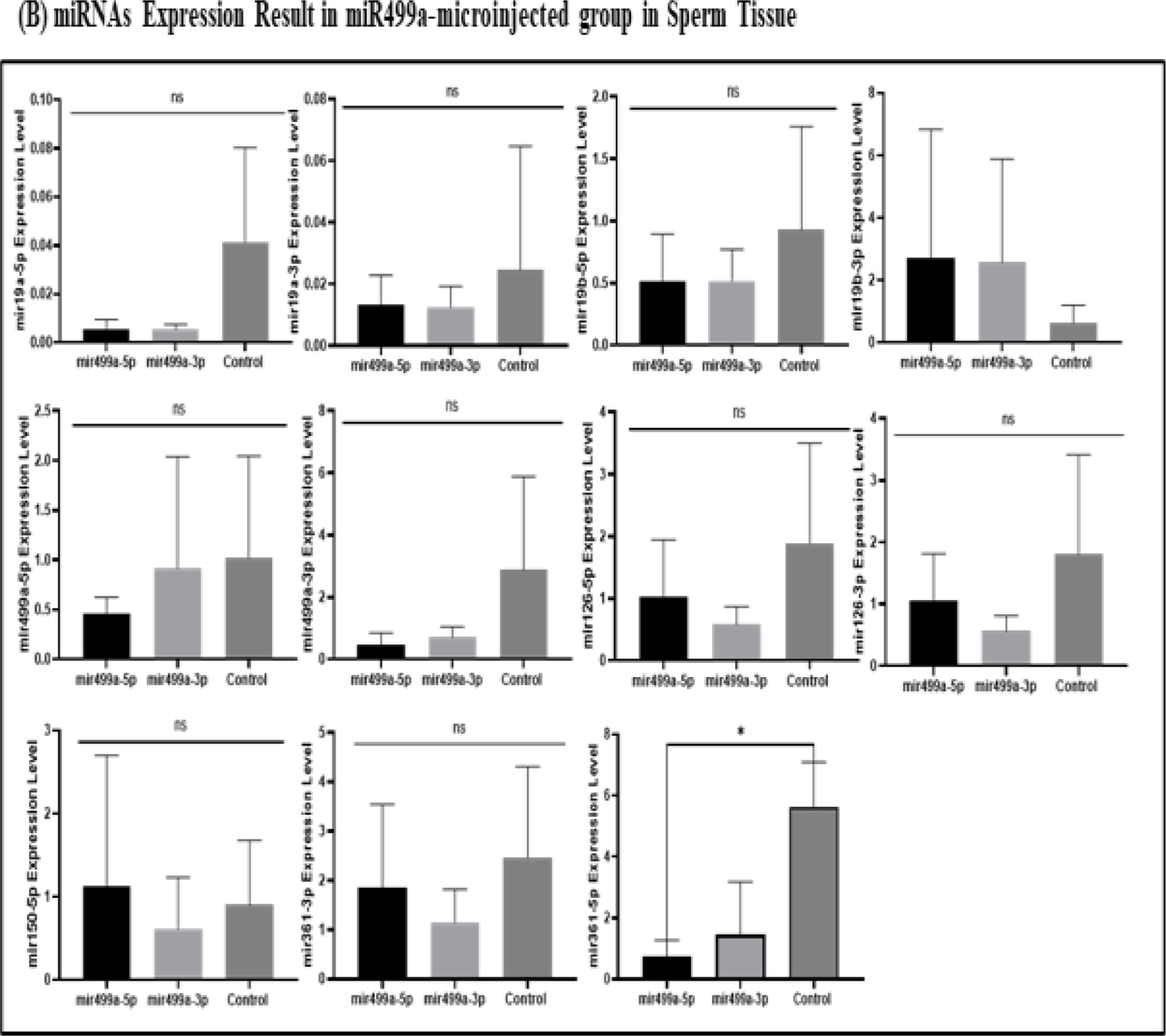
miRNAs’ expression results in sperm. (**A**) miRNAs’ expression results in miR-19a microinjected group (5p/3p), (**B**) miRNAs’ expression results in miR-499a microinjected group (5p/3p).

### DNA-bound miRNA varies with changes in its own levels

In an abstract graphical summary, Figure 9 (blood) and Figure 10 (sperm) show the altered levels of the six miRNAs (in free fraction). Next, we investigated whether the decay of miRNAs in blood cells is also altered in the fraction of miRNAs bound to DNA? Because, we have previously reported that part of the nascent transcripts are retained on the genomic DNA in the hybrid structure. To extract the total RNAs retained on the genome see the Method section, briefly, the genomic DNA is first purified from the cells then the RNAs retained on the R-Loop hybrid structure (DNA/RNA) are extracted by an extensive treatment with *Dnase*. As expected, a strong reduction in miRNAs retained on the genome is observed in tissues samples from animals born after microinjection of miRNA. However, the decay of six miRNAs was tissue-specific (blood, hippocampus and sperm), for example the decay of miR-19a in blood samples was highly specific for miR-19a-5p with microinjection of miR-19a-5-p and at a significantly higher or unchanged level with miR-19a-3p microinjection (Figure 11, 12 and 13). In contrast, decay in the hippocampus was specific for miR-499a-5p with miR-499a-5p microinjection and greater or unchanged with miR-499a-3p microinjection (Figure 14, 15 and 16 with graphical abstract Figure 17).

**Figure 9:**
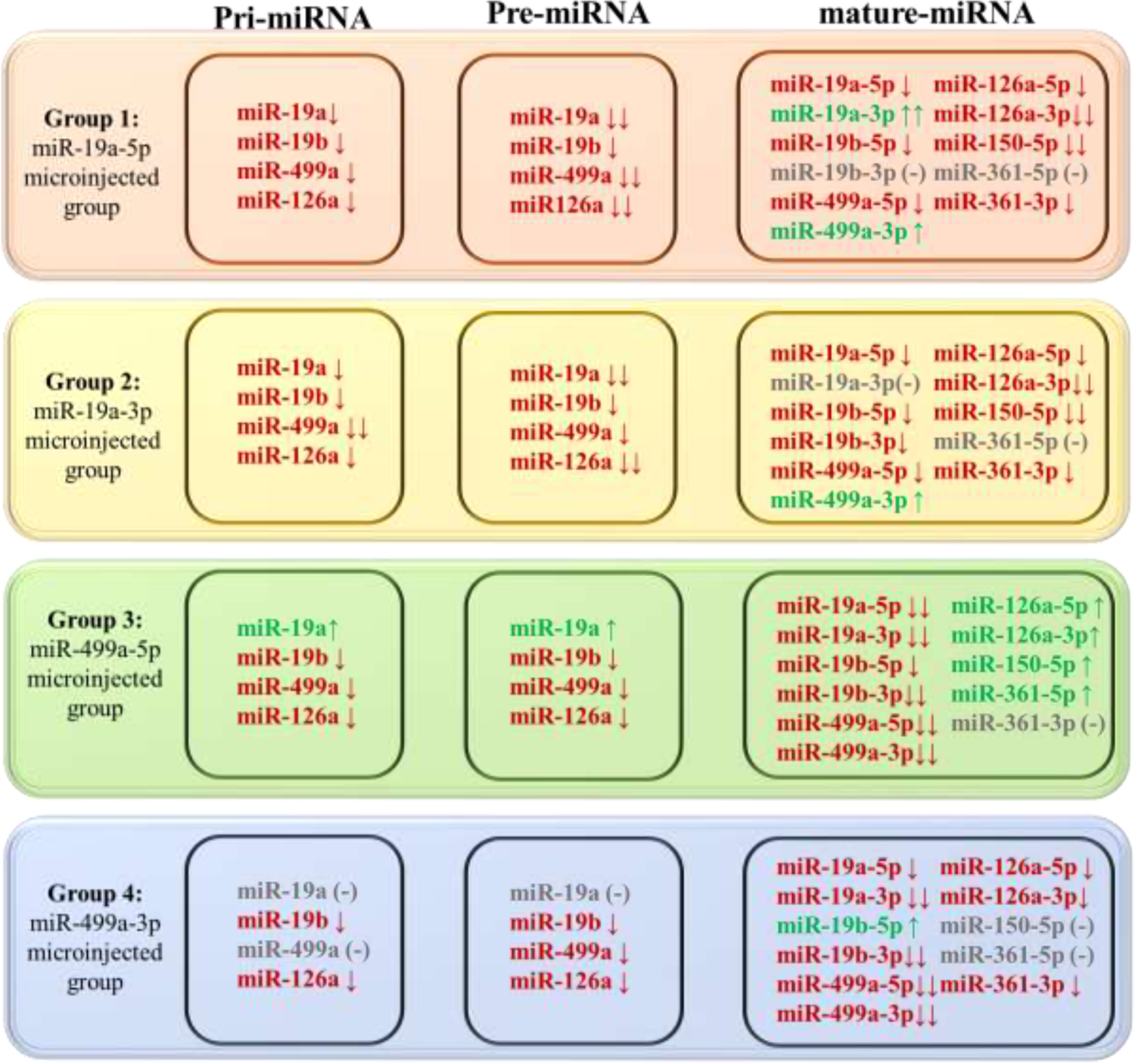
Graphical abstract of blood pri-pre and mature miRNA expression level changes comparing with control group. (Red arrow: down-regulation; green arrow: up-regulation; (-) not changing.)

**Figure 10:**
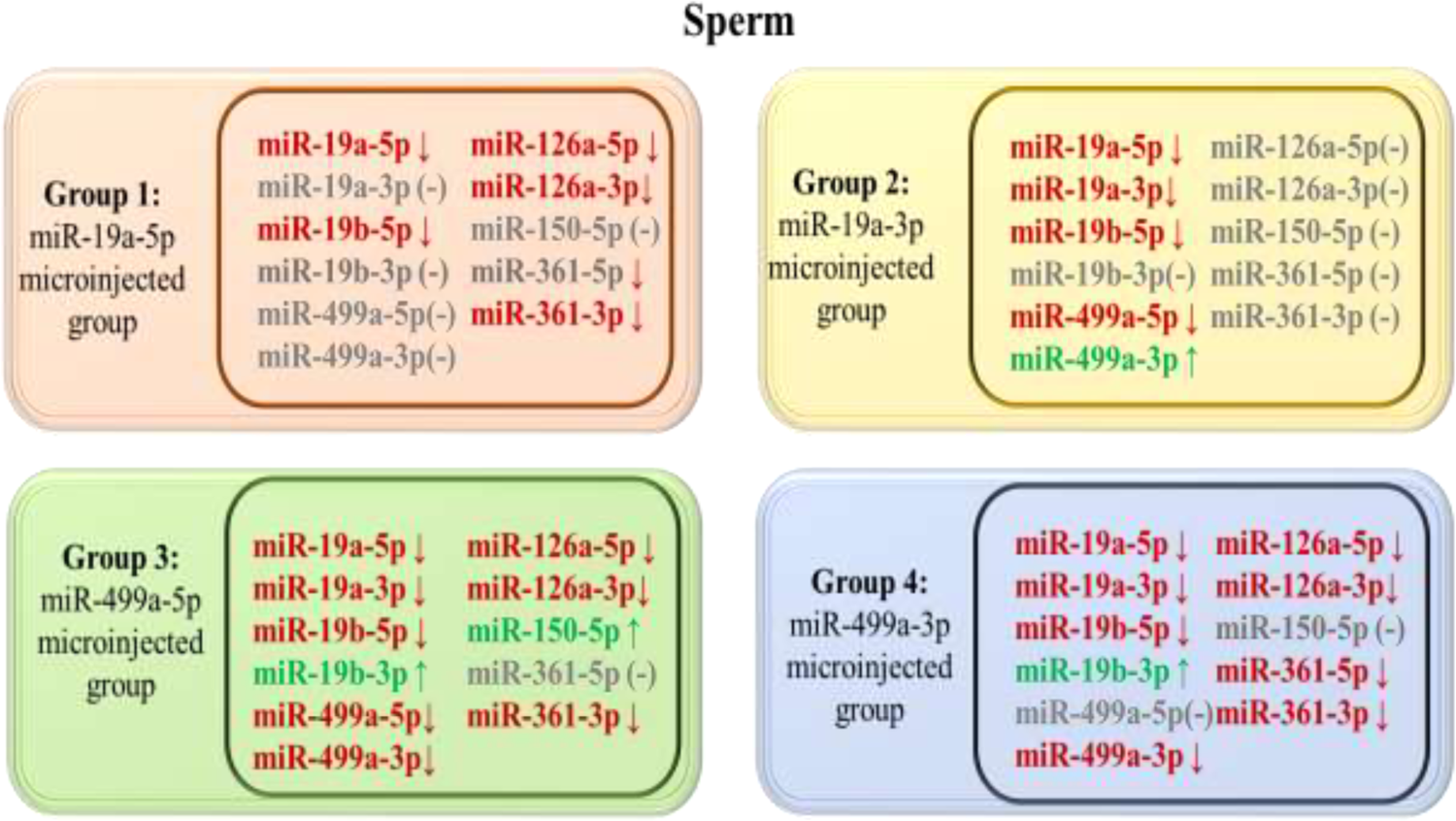
Graphical abstract of sperm pri-pre and mature miRNA expression level changes compared with the control group. (Red arrow: down-regulation; green arrow: up-regulation; (-) not changing).

**Figure 11:**
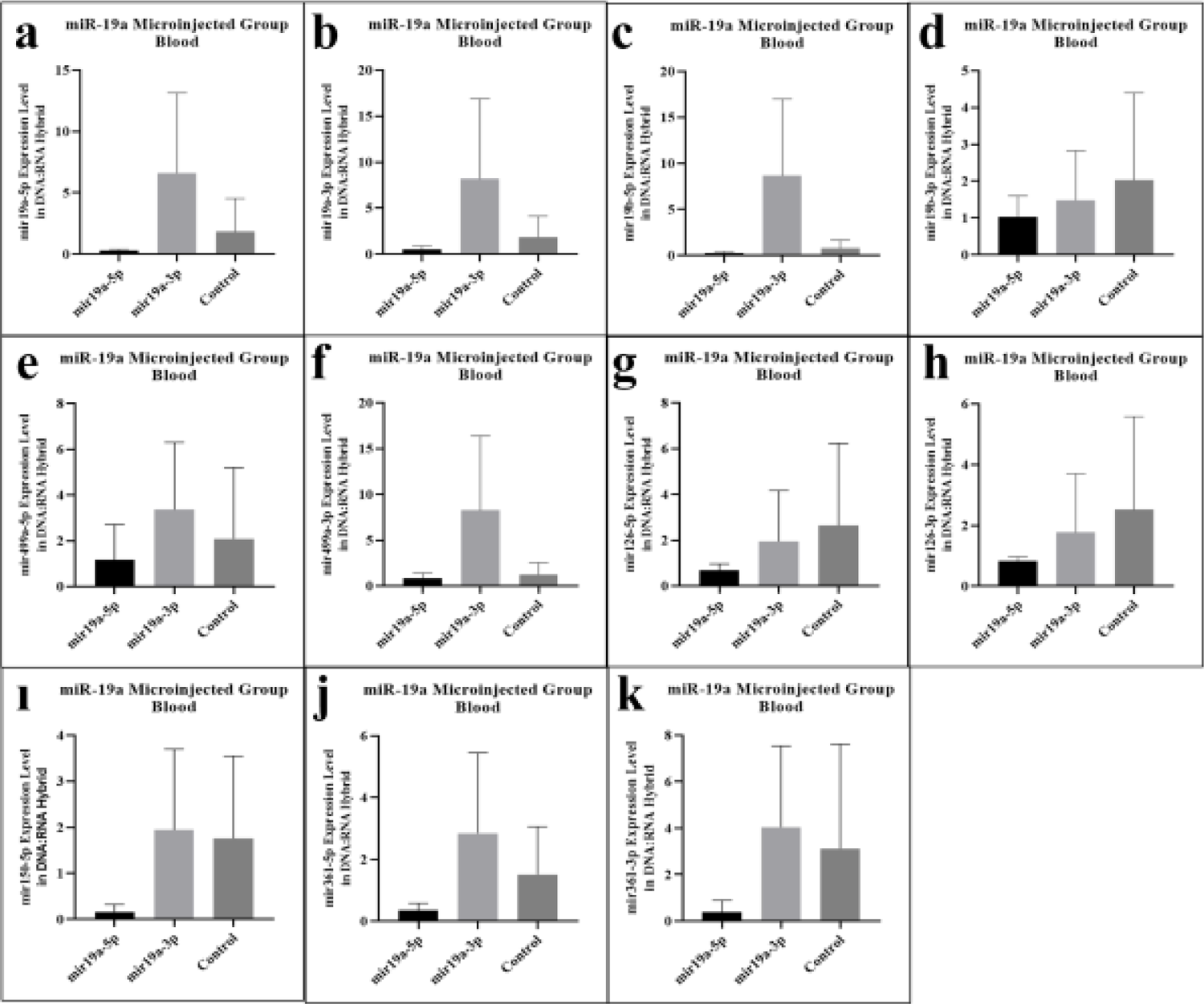
miRNAs expression results in blood DNA:RNA hybrid in miR-19a microinjected mice. **a.** miR-19a-5p expression level, **b.** miR-19a-3p expression level, **c.** miR-19b-5p expression level, **d.** miR-19b-p expression level, **e.** miR-499a-5p expression level, **f.** miR-499a-3p expression level, **g.** miR-126a-5p expression level, **h.** miR-126a-3p expression level, **ı.** miR-150a-5p expression level, **j.** miR-361a-5p expression level and **k.** miR-361a-3p expression level.

**Figure 12:**
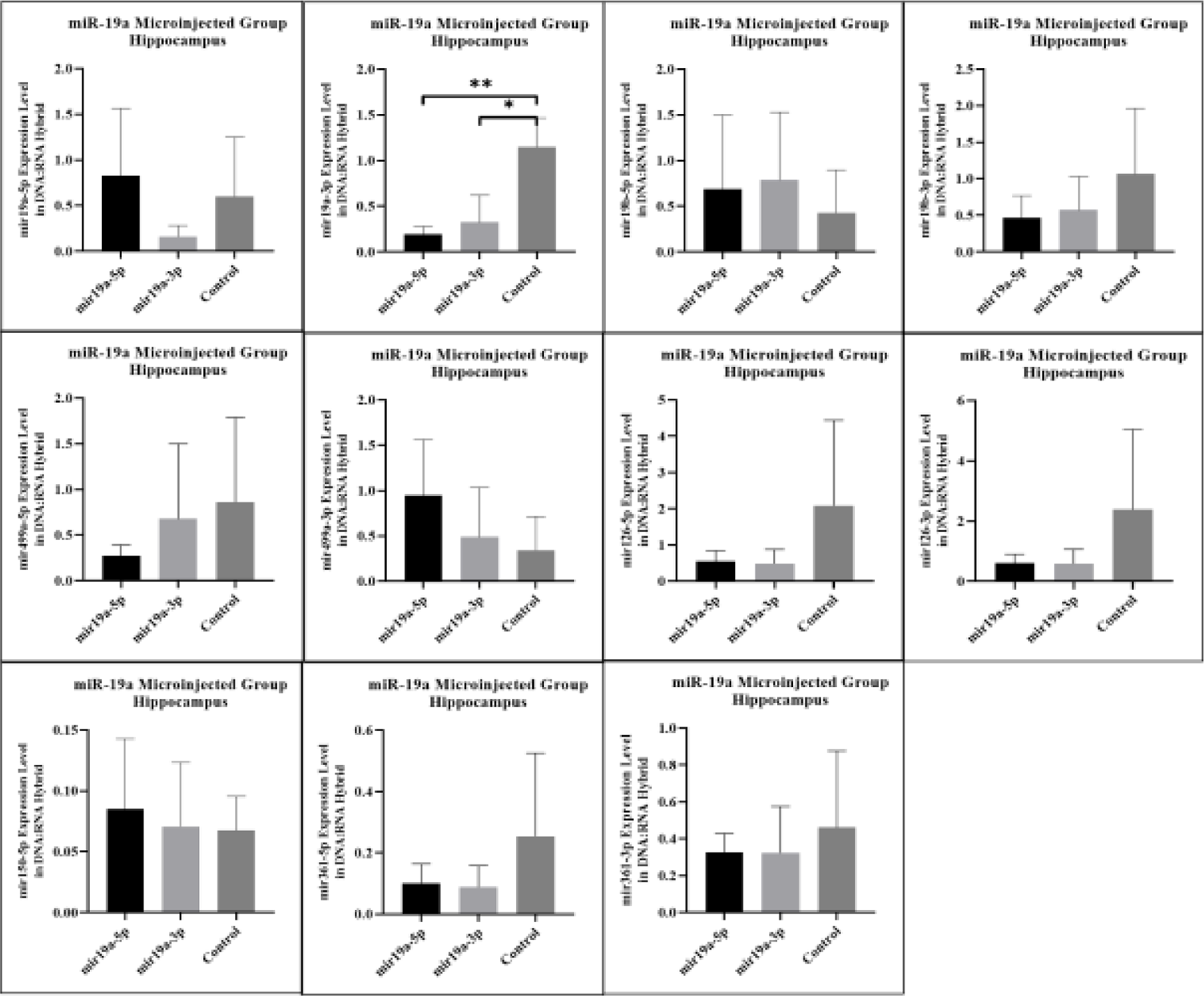
miRNAs expression results in hippocampus DNA:RNA hybrid in miR-19a microinjected mice. **a.** miR-19a-5p expression level, **b.** miR-19a-3p expression level, **c.** miR-19b-5p expression level, **d.** miR-19b-p expression level, **e.** miR-499a-5p expression level, **f.** miR-499a-3p expression level, **g.** miR-126a-5p expression level, **h.** miR-126a-3p expression level, **ı.** miR-150a-5p expression level, **j.** miR-361a-5p expression level and **k.** miR-361a-3p expression level.

**Figure 13:**
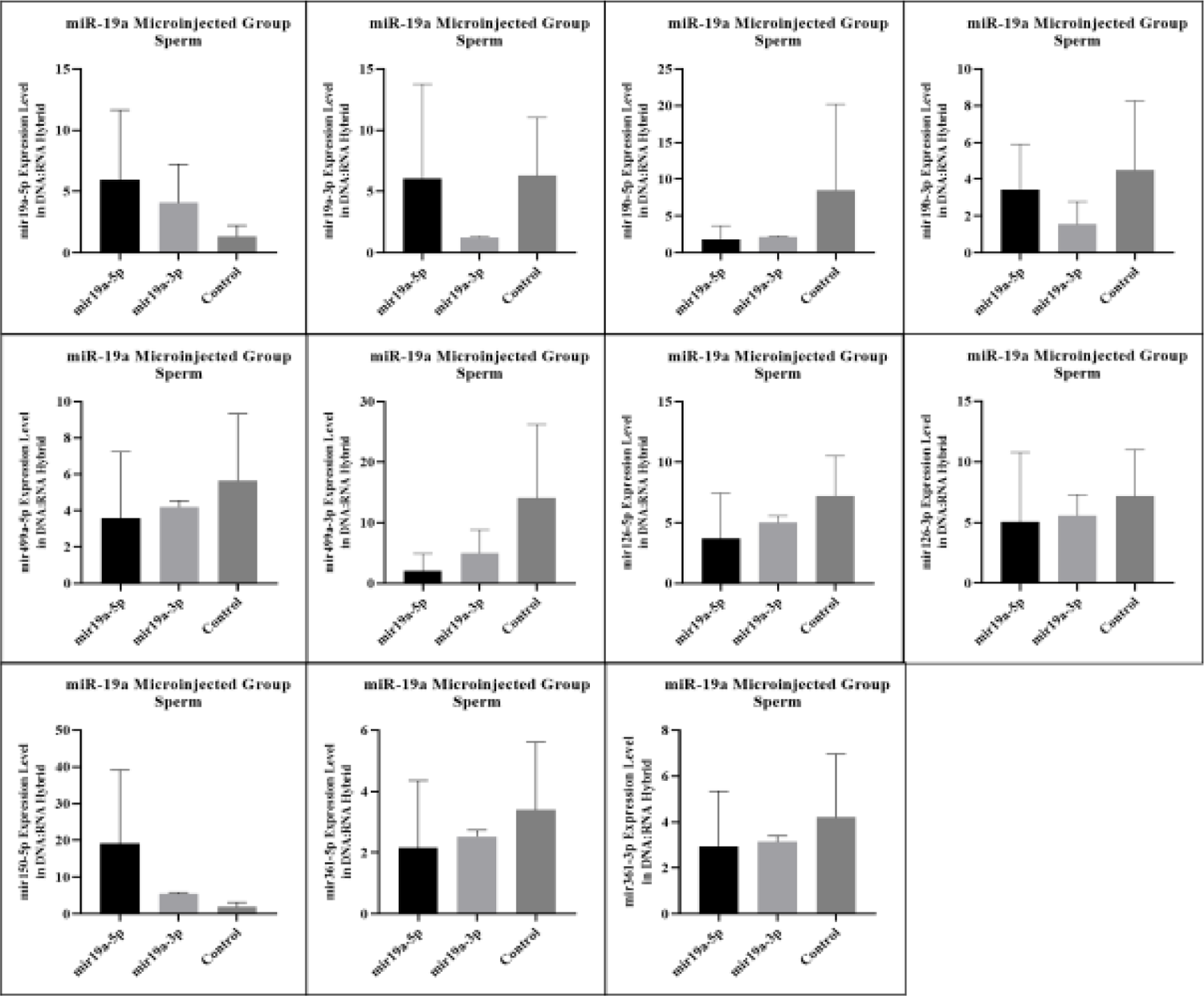
miRNAs expression results in sperm DNA:RNA hybrid in miR-19a microinjected mice. **a.** miR-19a-5p expression level, **b.** miR-19a-3p expression level, **c.** miR-19b-5p expression level, **d.** miR-19b-p expression level, **e.** miR-499a-5p expression level, **f.** miR-499a-3p expression level, **g.** miR-126a-5p expression level, **h.** miR-126a-3p expression level, **ı.** miR-150a-5p expression level, **j.** miR-361a-5p expression level and **k.** miR-361a-3p expression level.

**Figure 14:**
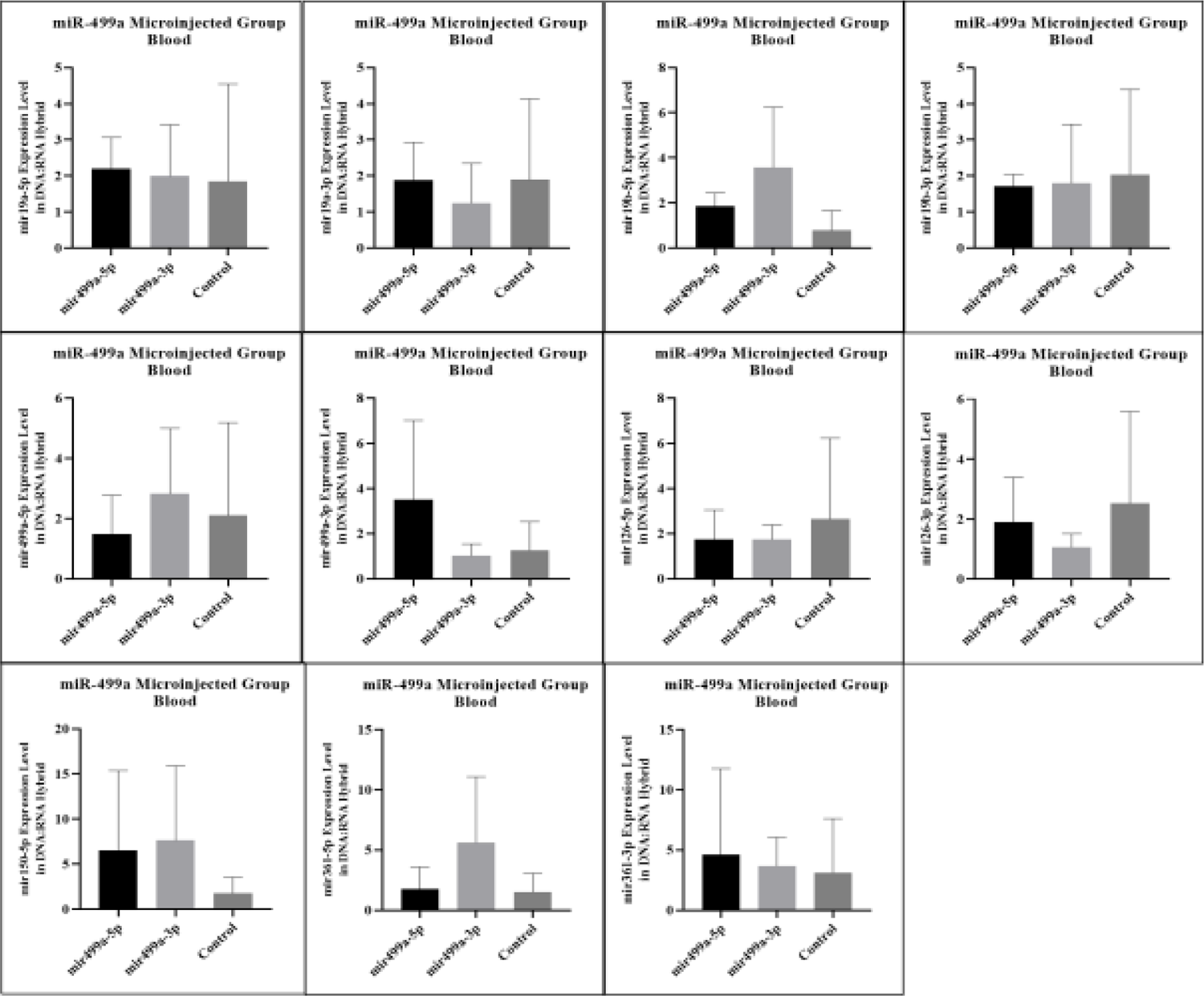
miRNAs expression results in blood DNA:RNA hybrid in miR-499a microinjected mice. **a.** miR-19a-5p expression level, **b.** miR-19a-3p expression level, **c.** miR-19b-5p expression level, **d.** miR-19b-p expression level, **e.** miR-499a-5p expression level, **f.** miR-499a-3p expression level, **g.** miR-126a-5p expression level, **h.** miR-126a-3p expression level, **ı.** miR-150a-5p expression level, **j.** miR-361a-5p expression level and **k.** miR-361a-3p expression level.

**Figure 15:**
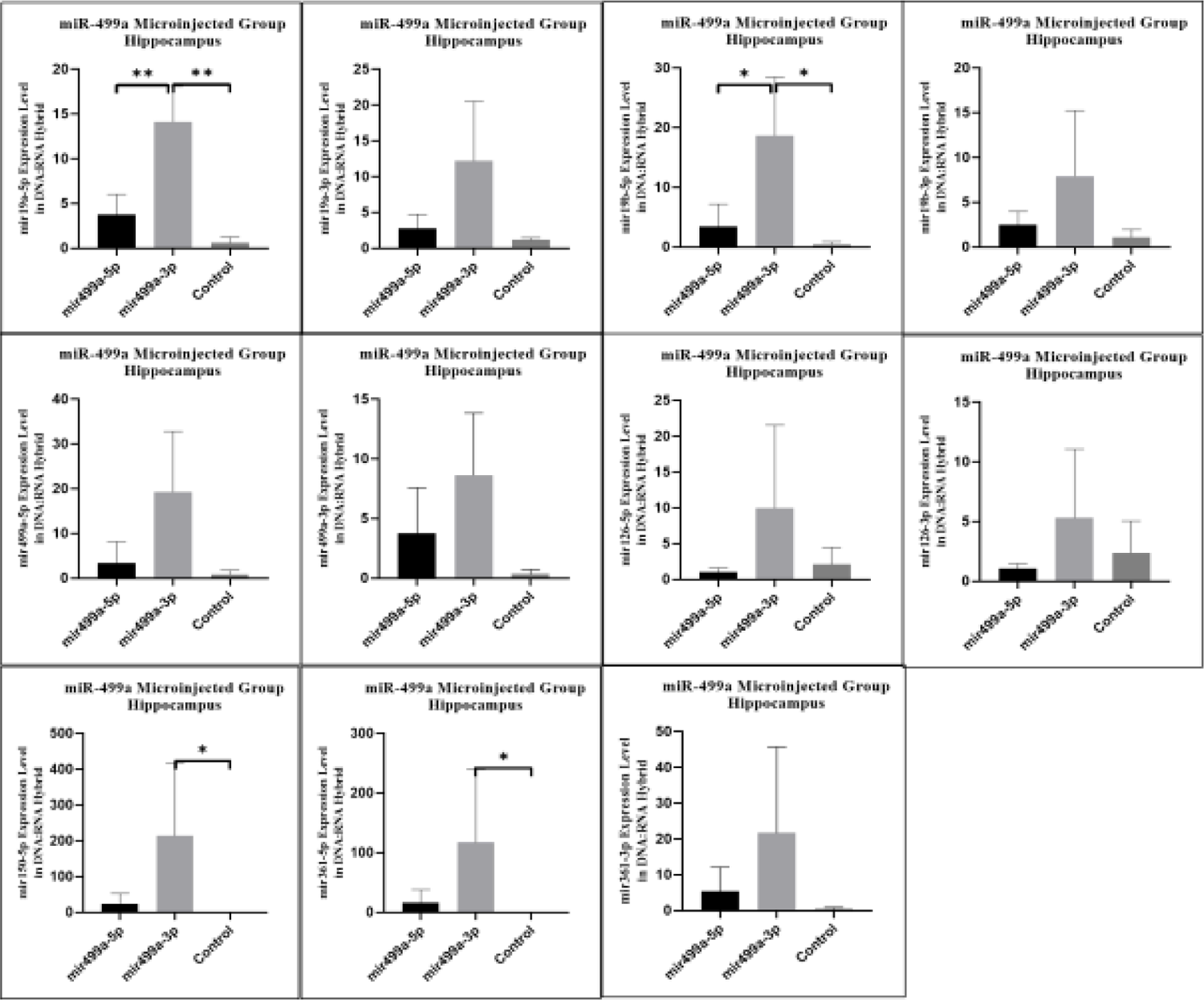
miRNAs expression results in hippocampus DNA:RNA hybrid in miR-499a microinjected mice. **a.** miR-19a-5p expression level, **b.** miR-19a-3p expression level, **c.** miR-19b-5p expression level, **d.** miR-19b-p expression level, **e.** miR-499a-5p expression level, **f.** miR-499a-3p expression level, **g.** miR-126a-5p expression level, **h.** miR-126a-3p expression level, **ı.** miR-150a-5p expression level, **j.** miR-361a-5p expression level and **k.** miR-361a-3p expression level.

**Figure 16:**
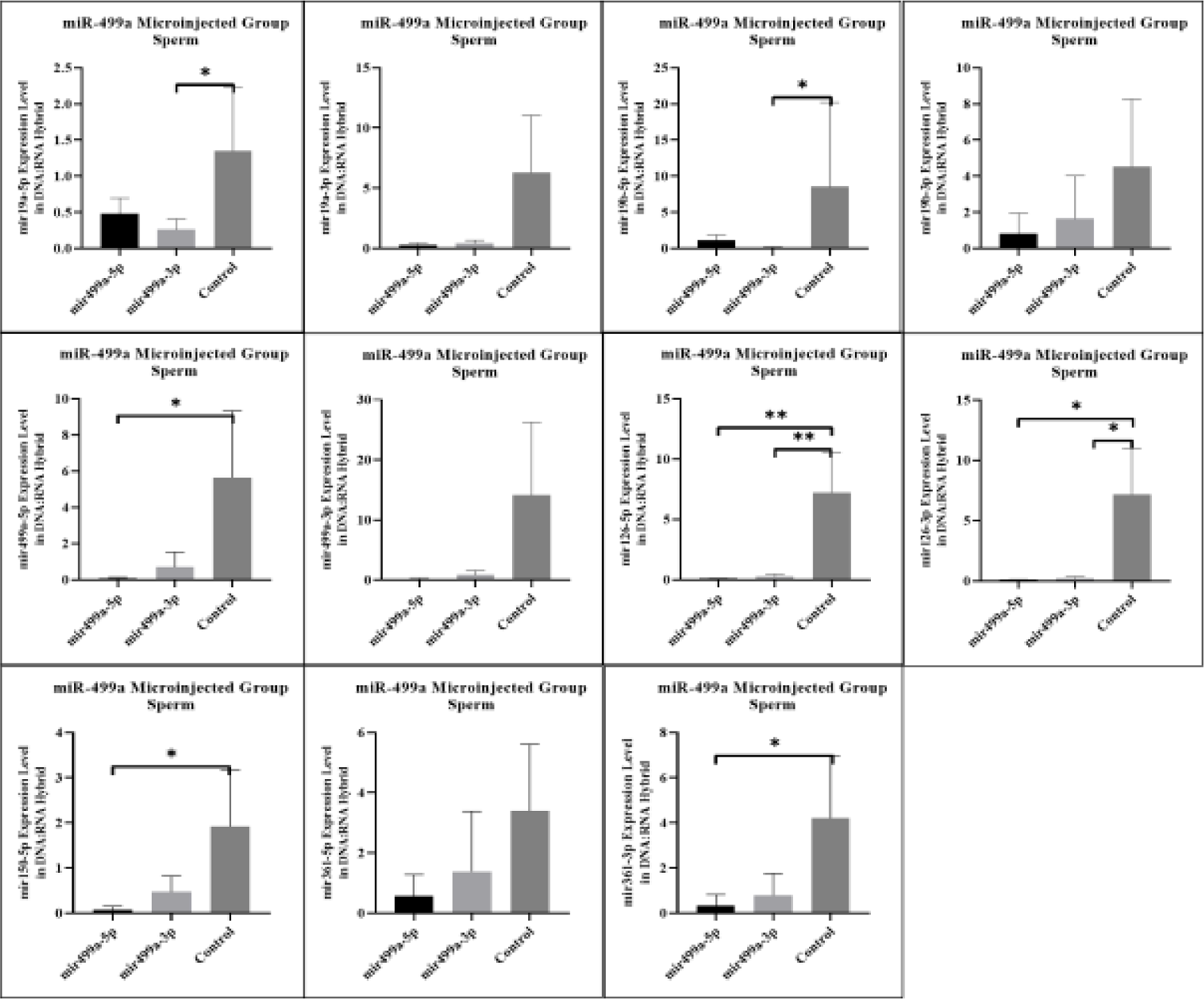
miRNAs expression results in sperm DNA:RNA hybrid in miR-499a microinjected mice. **a.** miR-19a-5p expression level, **b.** miR-19a-3p expression level, **c.** miR-19b-5p expression level, **d.** miR-19b-p expression level, **e.** miR-499a-5p expression level, **f.** miR-499a-3p expression level, **g.** miR-126a-5p expression level, **h.** miR-126a-3p expression level, **ı.** miR-150a-5p expression level, **j.** miR-361a-5p expression level and **k.** miR-361a-3p expression level.

**Figure 17:**
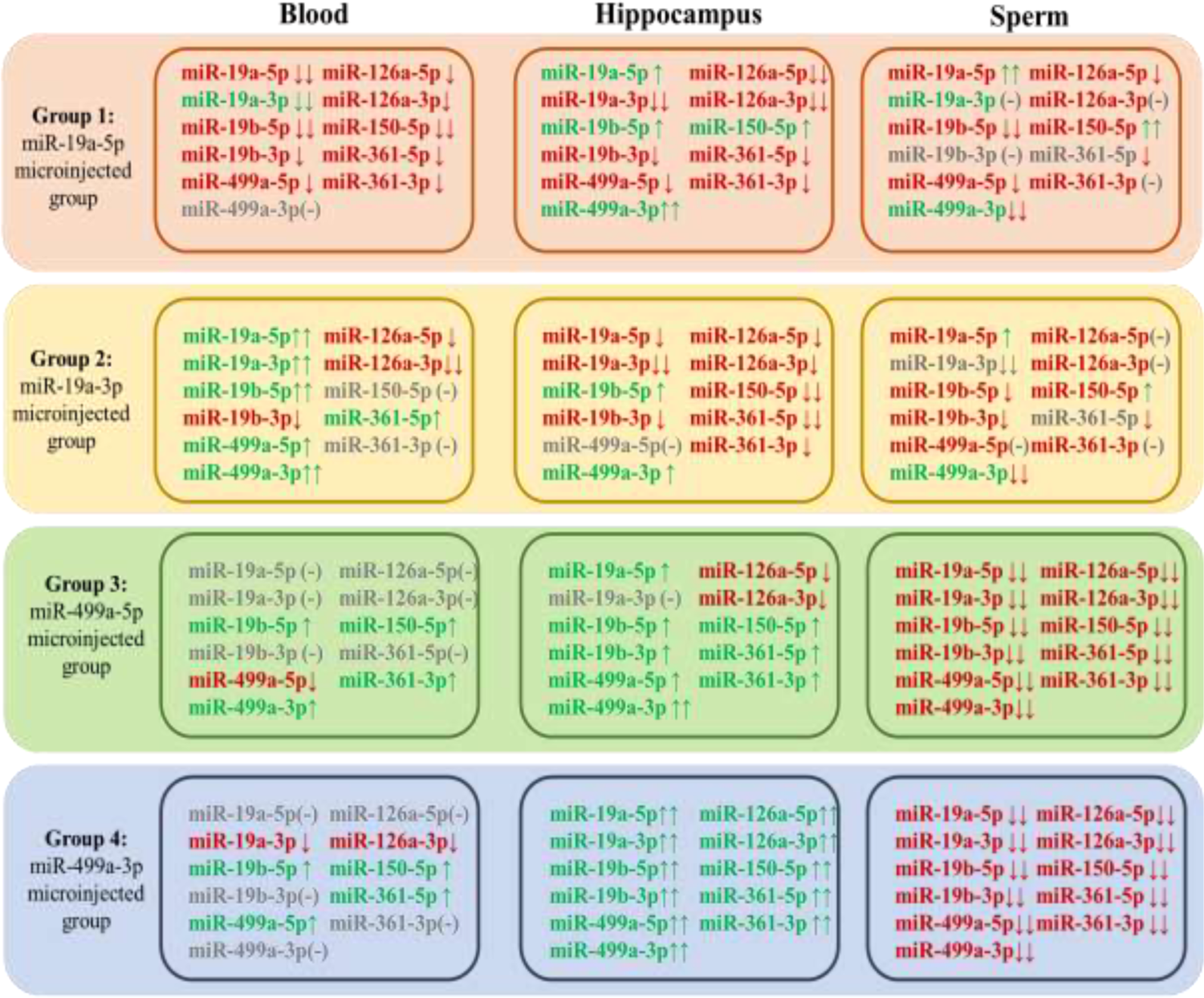
Graphical abstract of mature miRNA expression level changes in DNA: RNA hybrid comparing with control group in blood, hippocampus and sperm tissue. (Red arrow: down-regulation; green arrow: up-regulation; (-) not changing.)

To confirm and extend these results to human samples, we performed the same DNA-bound RNA extraction protocol from the blood cells of six patients or six controls. All six miRNAs were also at lower levels in the DNA-bound RNA fraction of patient samples compared to controls Figure 18. Consistent with the animals results, most of the six miRNAs showed a strong decrease and two (miR-499a-3p and miR-19b-5p) resisted to this decrease. The change in distribution of other members of the six miRNAs was also observed. The overall analysis further demonstrated that the decay selectivity of the six miRNAs (DNA-bound RNA) in blood represent the six most highly down-regulated miRNA in these patient samples. In contrast, in patient blood samples miR-499a-3p remains elevated relative to the control. This analysis reiterates the tissue specificity and effect of 5p versus 3p miRNA sequences in the DNA-bound miRNA fraction (Figure 18, with graphical abstract Figure 19).

**Figure 18:**
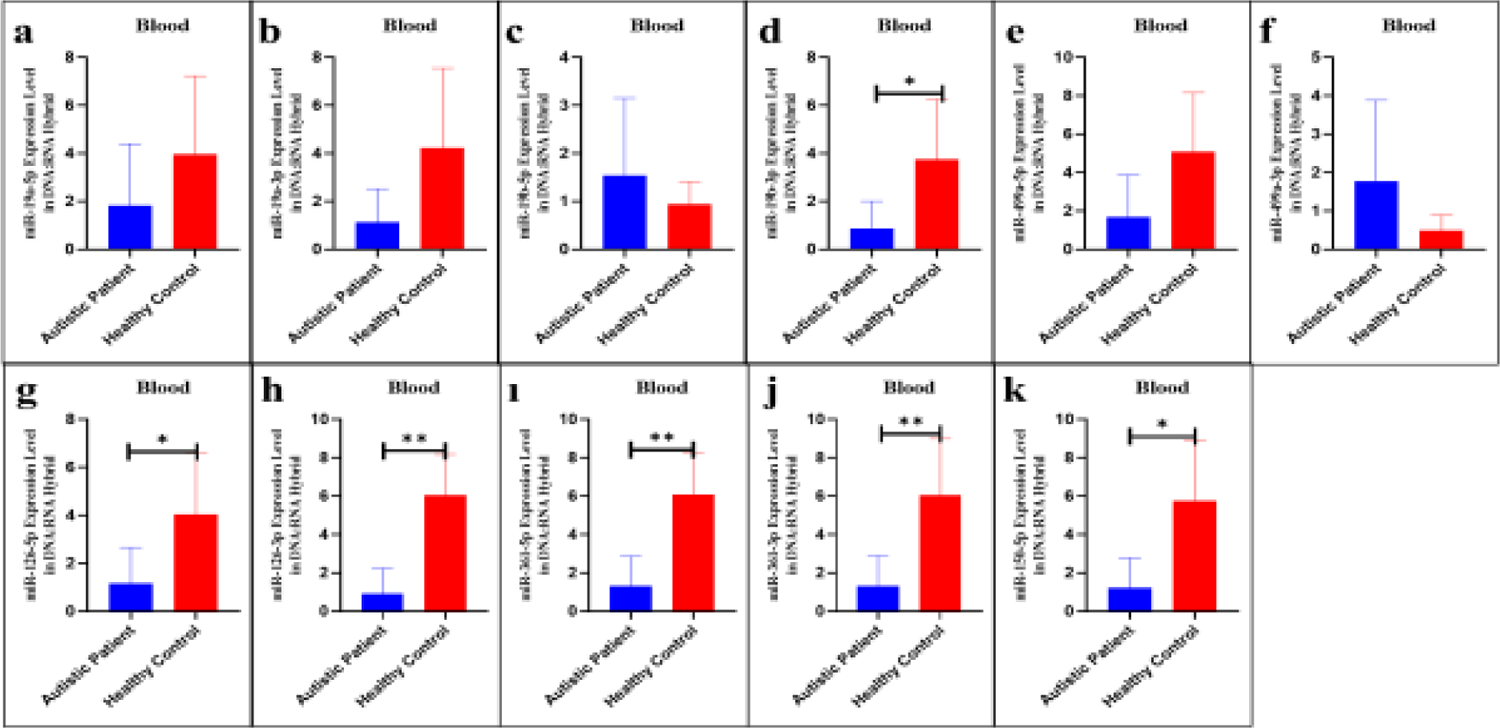
miRNAs expression results in blood DNA:RNA hybrid in human autistic patient. **a.** miR-19a-5p expression level, **b.** miR-19a-3p expression level, **c.** miR-19b-5p expression level, **d.** miR-19b-p expression level, **e.** miR-499a-5p expression level, **f.** miR-499a-3p expression level, **g.** miR-126a-5p expression level, **h.** miR-126a-3p expression level, **ı.** miR-150a-5p expression level, **j.** miR-361a-5p expression level and **k.** miR-361a-3p expression level.

**Figure 19:**
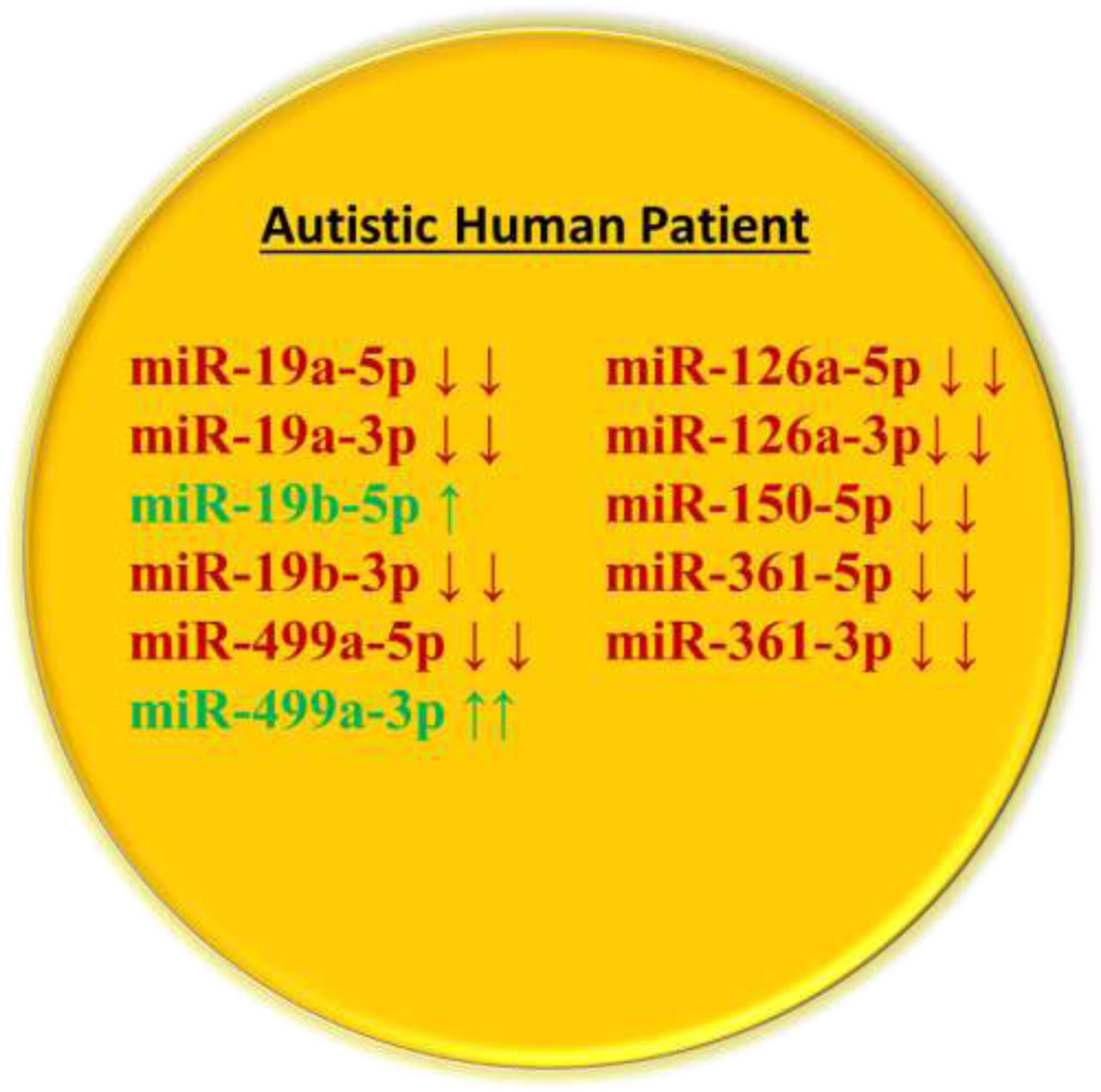
Graphical abstract of mature miRNA expression level changes in DNA:RNA hybrid comparing with control group in blood tissue. (Red arrow: down-regulation; green arrow: up-regulation; (-) not changing.)

Together, these results show that the decay of the six miRNAs at the DNA/RNA hybrid level is a selectively regulated decay process, rather than simply the result of synchronous decay through the post-transcriptional pathway, since miRNAs levels derivatives of the same transcript can be decoupled. Moreover, these results show that the sequence of the six miRNAs is necessary for this regulated decay.

## Discussion

Despite overwhelming evidence for the level of transcription of miRNAs in normal development, the mechanism by which changes in the level (proportion) of these regulatory transcripts could eventually increase the risk of pathologies is not understood. The aim of this study was to down-regulate specific miRNAs in serum and to assess phenotypic and transcriptional changes in mice. We hypothesized that quantitatively changing the miRNAs quota level of fertilized mouse eggs will impact developing tissues. Now, we provide a direct and unique method to modify the initial levels of a miRNA and its associated pathways.

All results on miRNAs expression levels have been summarized in Figures 9, 10, and 17 (mouse) and 19 (human) as a graphical abstract. Upon microinjection of each miRNA into fertilized mouse eggs, in this study, analysis of miRNA expression profiles in serum, hippocampus and sperm revealed that six miRNAs were altered in accordance with behavior changes. We confirmed that expression of serum miRNAs level was lower in mouse models with affected behavior than in healthy controls. Interestingly, the regulation of miRNA expression was found to be dependent on the type of miRNAs within the tested group which also showed transcriptional alterations already visible at their pri- and pre-miRNAs. Each of the miRNA among the test groups while microinjected into fertilized mouse eggs induces a specific phenotype and miRNAs profiles. Our results indicate that specific miRNAs could be potential regulators of their own transcriptions.

One mechanism could be that variation in miRNAs level in fertilized eggs affects gene dosage in embryos and is therefore maintained into adulthood via a transcriptional cascade. That is, the original miRNA pool, which should target the main regulatory transcription networks of developing embryos, and then the following chain reactions guide to reprogram variable epigenetic profiles in different tissues including the blood cells of the offspring. Changes in the miRNAs pool affect several organ development pathways^14–16^. Both genetic and epigenetic alterations have been shown to cause abnormal expression of miRNAs (see for review ^17^). This negative response after a large amount of miRNAs microinjected into eggs, is reminiscent of Paramutation events in the plant that depend on a threshold of small RNAs for trans-communication to establish and maintain chromatin states^18^. Moreover, in these plants the expression levels from B’ and BI loci depend on *mop1* (RDR) necessary to maintain this threshold. In the absence of RDR in mouse how these thresholds are maintained, is still very mysterious.

Additionally, of the six miRNAs, miR-19 has been reported in at least three separate reports to be impaired in autistic patients^19–21^. In particular, miR-19 present in miR-17-92a clusters a well-known miRNAs cluster, has been shown to vary with changes in environmental factors ^17, 22, 23^. The results to date lead to show that the expression levels of microRNAs or a certain number of them can vary according to environmental changes^24^.

These results support our hypothesis that miRNAs respond to quantity changes by modifying their own transcription. Here, fertilized eggs overloaded with miRNAs down-regulated their transcription, which may alter the amounts of other miRNAs. We showed here that lower expression levels of each miRNA (miR-19 or miR-499) were associated with specific behavioral changes. Indeed, in our data set, we found that changes in animal group serum miRNAs levels were associated with the recurrence of behavior changes. Each microRNAs contributes in part to the phenotypic changes of a complex disease such as autism. A full autism phenotype would likely require down-regulation of at least so far all six miRNAs, as down-regulation of each miRNA only affects a given phenotype. In human analysis of miRNAs distributions in patients and siblings, this also suggests that down regulation of all six miRNAs is necessary in an child with autism.

### miRNAs levels participate in the expression of the phenotype

Recently, Ozkul et al.^5^ reported that a panel of five miRNAs (miR-19a-3p, miR-361-3p, miR-150-5p, miR-126-3p and miR-499a-5p) are down-regulated after VPA treatment or genetic modifications (*Cc2d1+/-*) in mice. These data were consistent with our results that these miRNAs were significantly downregulated in the serum of mouse models, and our current data even provided evidence that the down-regulation of two of these miRNAs separately in the serum was accompanied by certain behavior changes. miRNAs are deregulated in adult tissues when their fertilized eggs have been exposed to a high level of miRNAs. These results suggest that miRNA levels are regulated at an early stage of development.

We have observed a relationship between the level of specific miRNAs (19a-, 499a-) and phenotype changes, but the mechanisms by which the specific altered miRNAs act are unknown in detail. Post-transcriptional regulatory mechanisms at the levels of individual transcripts are precisely detailed, but the orchestration of genome-wide transcription is still in obscurity with the possibility that the enormous amount of noncoding RNAs y participate. Several reports have shown that miRNAs are usually transcribed by RNA polymerase-II, a clarification of the signaling pathway upstream of miRNAs is definitely needed.

### Could DNA-bound miRNAs control their own transcripts?

The organization of nuclear compartments is proposed as also dependent on non-coding RNAs (ncRNAs) ^25, 26^. A number of questions could then be raised. It is not yet clear how subtle changes in miRNAs levels might influence these nuclear compartments? Elevated/decreased levels of a given miRNAs could be factored into overall changes in RNA-RNA, RNA-DNA and DNA/DNA interactions in the nucleus, but how are they stored and propagated for transfer to daughter cells or the next generation still uncertain.

Our data suggest that the regulatory mechanisms of the tested miRNA*s* differ from the cytoplasmic or TDMD post-transcription pathways previously described. First, decay is inherited from an altered stage of one cell embryo as cellular memory. Second, the decrease is also observed in the fraction of the last wave of transcriptional base-pairing with its own trigger DNA. Together, these data suggest that at least two of these six-miRNAs are initially regulated by their own quantity and DNA/RNA hybrid sequence binding in addition to their role in the classical seed-dependent post-transcriptional mechanism, TDMD, and may – be a variant of it.

We propose a decay model of *six* miRNA*s* in which it suffices to vary the quantity of miRNAs attached to their own DNA.

### Hybrid DNA/RNA as effectors of transcription in eukaryotes

The complexity of gene expression does not yet allow us to ask simple questions such as:

What are the fastest and most effective minimal specific signals possible to enhance or block transcription? How to arrive at the simultaneous and specific control of the variation of the transcription of a group of genes in eukaryotes?

In eukaryotic cells, an individual promoter is used. This is how many transcripts (distributions of coding or non coding RNA) are produced which implement complex expression controls from the initiation of transcription to post-transcription. Although the transcriptional potency of promoters, enhancers and silencers is variable, it does not account for how eukaryotes might allow immediate and simultaneous transcriptional variation. RNA theory states that RNAs derived from different chromosomes act as specific functional information to generate their own transcriptional control and enable cohorts during simultaneous transcription. Simultaneous and efficient control of a gene transcription group is necessary for immediate co-transcription/silence of functionally related transcripts.

Why specific single-strand RNA as immediate transcriptional variation signals? This results exactly from the microinjection into fertilized mouse eggs of small synthetic RNAs oligonucleotides (21-22 ribo-nucleotides) see above therefore confirms our hypothesis. Indeed, following the microinjection of a large amount of RNA oligonucleotides, for example miR-19a-3p, into fertilized mouse eggs, the expression levels of the same miRNAs are down-regulated in the adult tissues. Microinjection of 5p or 3p strand of miRNA induces down-regulation of the same miRNAs. This negative regulation is at the transcriptional level, since pri-pre-miRNAs and fraction of nascent miRNAs levels are also reduced. The reduction in levels of RNAs attached to DNA suggests that RNA, through its homologous sequence and its interaction with DNA, regulates transcription levels. Here, we report the interaction of transcripts with the gene body. It is not yet clear if there is a specific required sequence length (here only 21 nucleotides) responsible for the regulatory impact? Or universally all produced RNAs are regulators of their own transcription. The multiplication of examples with complementary studies will demonstrate the precision with which the RNAs produced are their specific regulatory element for their own synthesis.

Work in mice has now strongly suggested this theory that simply RNAs could regulate their own synthesis and in this way the amount of RNA could influence transcription. We show here the down regulation artificially induced by an excess of miRNA but it is not yet clear, maybe the reverse effect how does upregulation occur? Moreover, it showed that associations of miRNAs with DNA create pathway-specific RNA expression that regulates the amount of miRNA with physiological impact. Importantly, we also observed down regulation of all six miRNAs as free miRNAs molecules and those attached to their own loci in patients with autism. Here, the involvement of the RNA/DNA hybrid in the formation and programmed expression of genes is proposed. The next important question is how to reverse, to get back to normal expression levels?

The DNA after being base paired with the miRNA sequence and eventually recruits an RNA-binding protein, which in turn can decrease/ or increase transcription. Alternatively, the miRNA could induce conformational changes in the hybrid structure (R-Loop) that directly alters transcription. If other possibilities exist, it will be with miRNAs closely linked to their DNA sequences. More experimental attempts to recognize miRNA sequences involved in transcription will further test the model of the sufficiency of miRNAs or seed sequence for this selective regulation.

Understanding the specific decay mechanism regulating transcription of miRNA will have an important impact. Such specific mechanisms are likely to be present in other biological systems because they allow specific and/ or simultaneous regulation of redundant RNA paralogs, allowing dynamic regulation of targets of an RNA family. In addition to functioning in normal physiology, sequence-specific decay mechanisms could provide an attractive avenue to modulate the abundance of specific miRNA families and their target genes. Transcript levels of mir-19 and miR-499 were altered; whether the strength of impact of these changes is similar in the context of all miRNAs or RNAs sequence will be a matter of future investigation. Having not examined processing intermediates (not easy in oocytes), we still have an incomplete understanding of how alterations introduced into a miRNA affect its synthesis. We did not directly measure the levels of miRNA decay at the embryonic stage, leaving ambiguity as to the molecular basis for the up or down regulation of miRNA variants in the embryo. Moreover, the decay of miR-19 occurs at a rate similar to that of miR-499; our interpretation is that these variants are subject to the same mechanism that targets DNA at its own locus. However, if the variants are targeted by a distinct decay mechanism within the same developmental time window, this could obscure the role in regulating the initial miRNA pool.

In conclusion, the regulation of a group of miRNAs in adult tissues depends on their quantity at an early stage of development. At this stage, it appears very mysterious compared to the known mechanisms. Further studies are needed to clarify the relationship between miRNAs and intercellular signaling in phenotype expression. These miRNAs could be a potential prognostic marker and a therapeutic target in behavior diseases.

### Figures legends

Mice born after microinjection of single stranded oligo RNA (microRNAs) into fertilized eggs, were designated miRNA* 5p end or 3p end see Table-1. In Figure 1 miRNAs level and behavior variations compared to those of controls.

## Supplementary supports

**Supplementary Figure 1:**
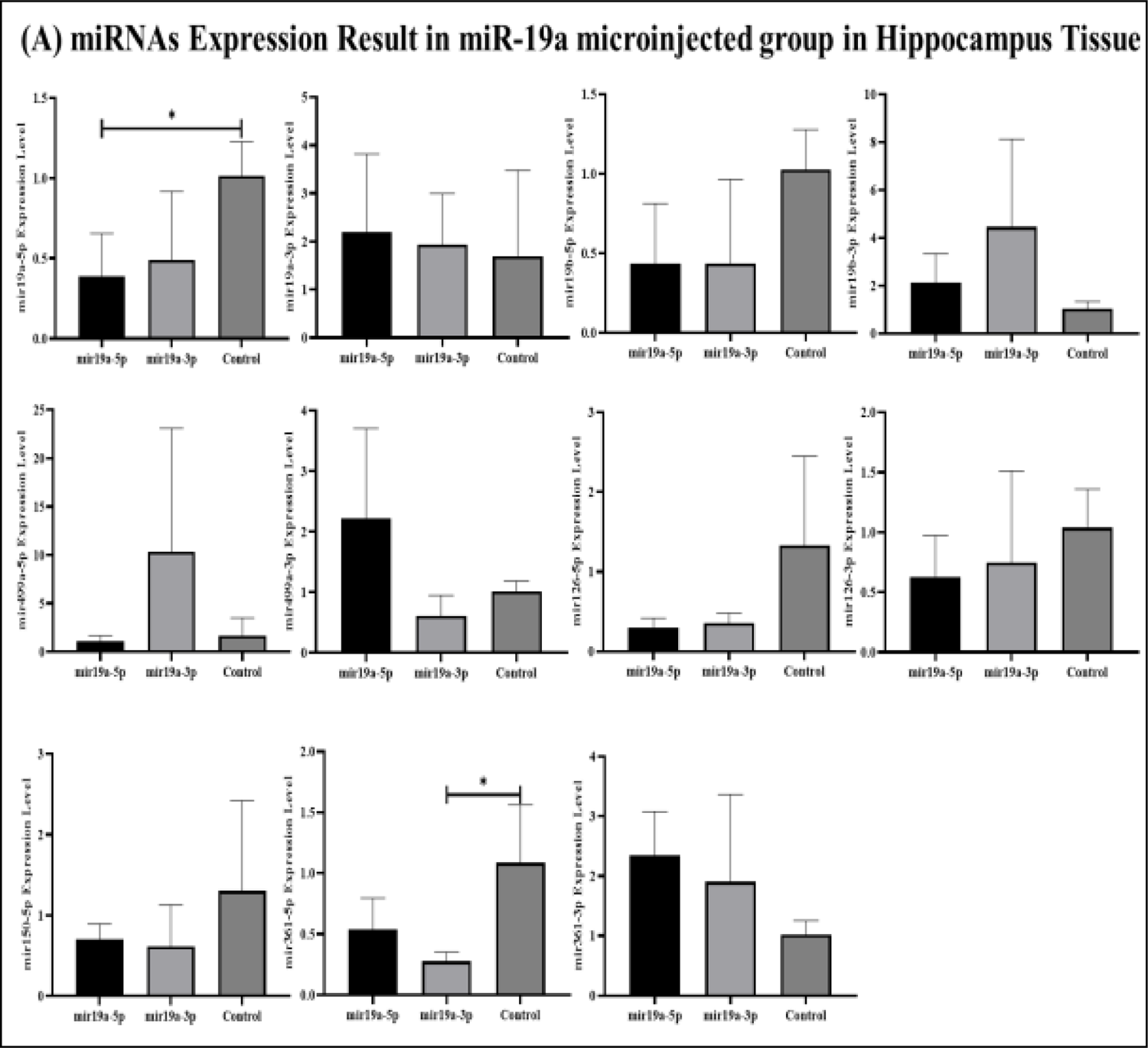
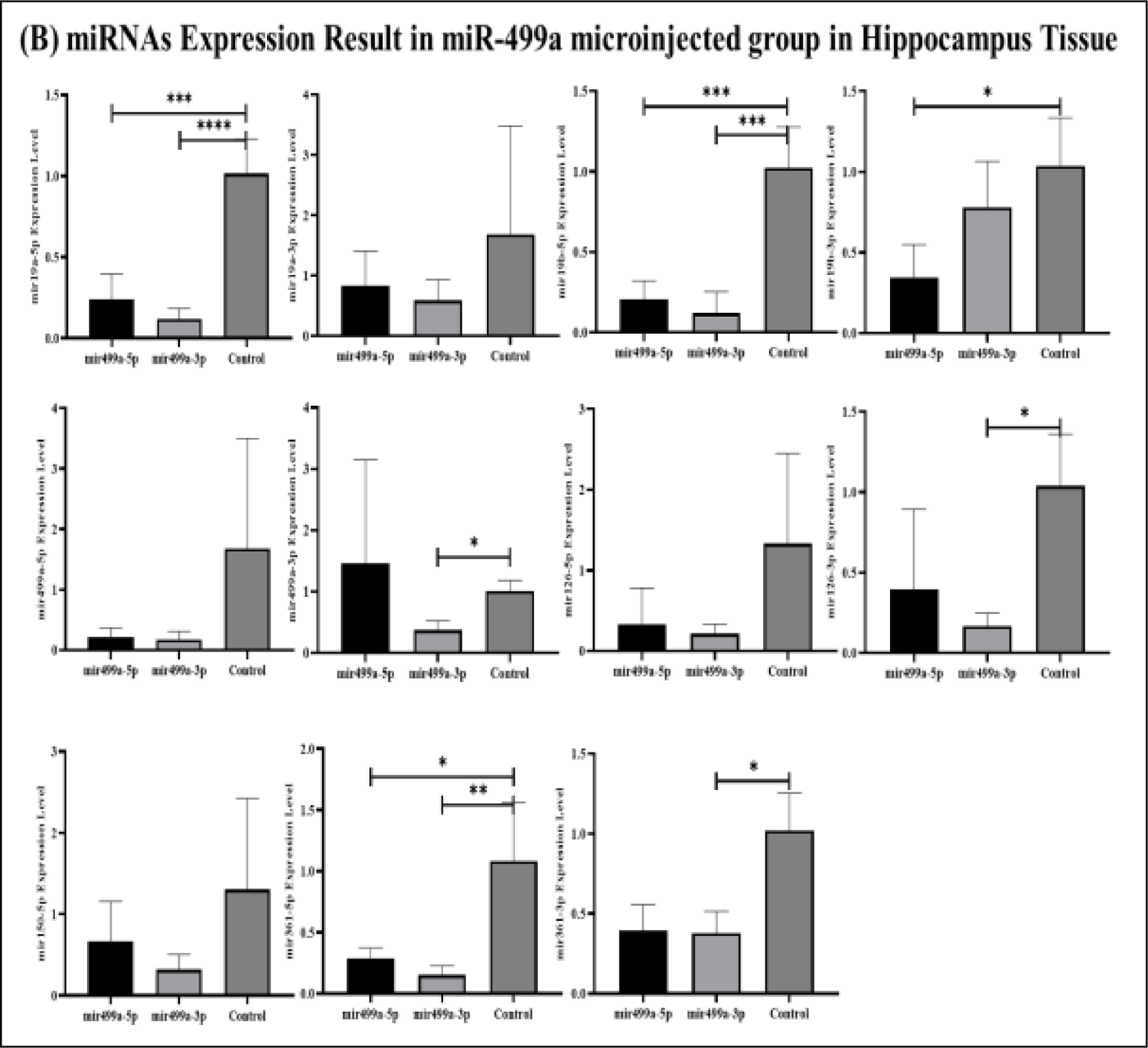
miRNAs expression results in the hippocampus. (A) miRNAs’ expression results in miR-19a microinjected group (5p/3p) (B) miRNAs’ expression results in miR-499a microinjected group (5p/3p).

**Supplementary Figure 2:**
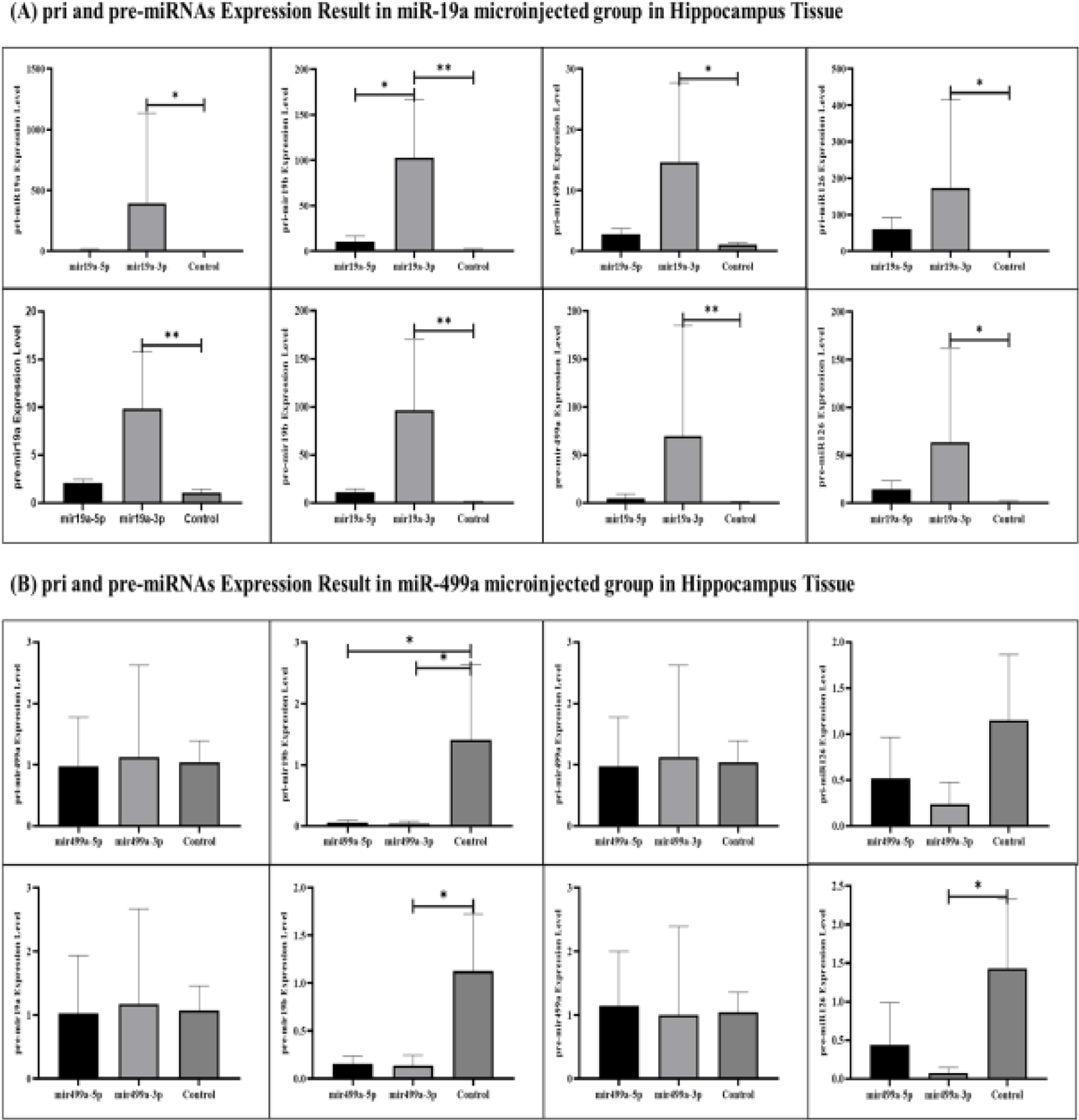
pri and pre-miRNAs’ expression results in the hippocampus. (A) pri and pre-miRNAs’ expression results in miR-19a microinjected group (5p/3p) (B) pri and pre-miRNAs’ expression results in miR-499a microinjected group (5p/3p).

**Table 1:**
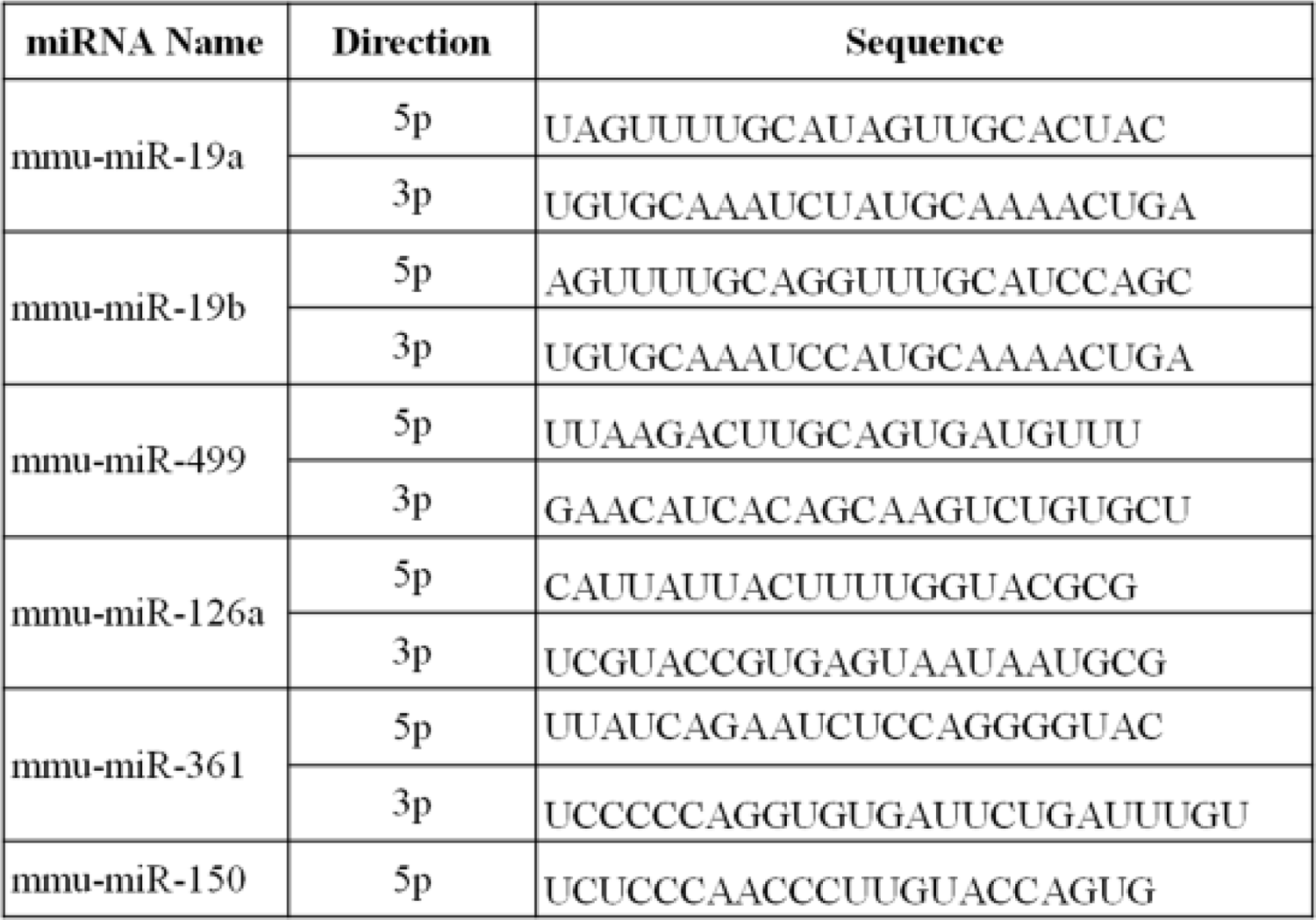
pre-miRNAs’ sequences.

**Table 2:**
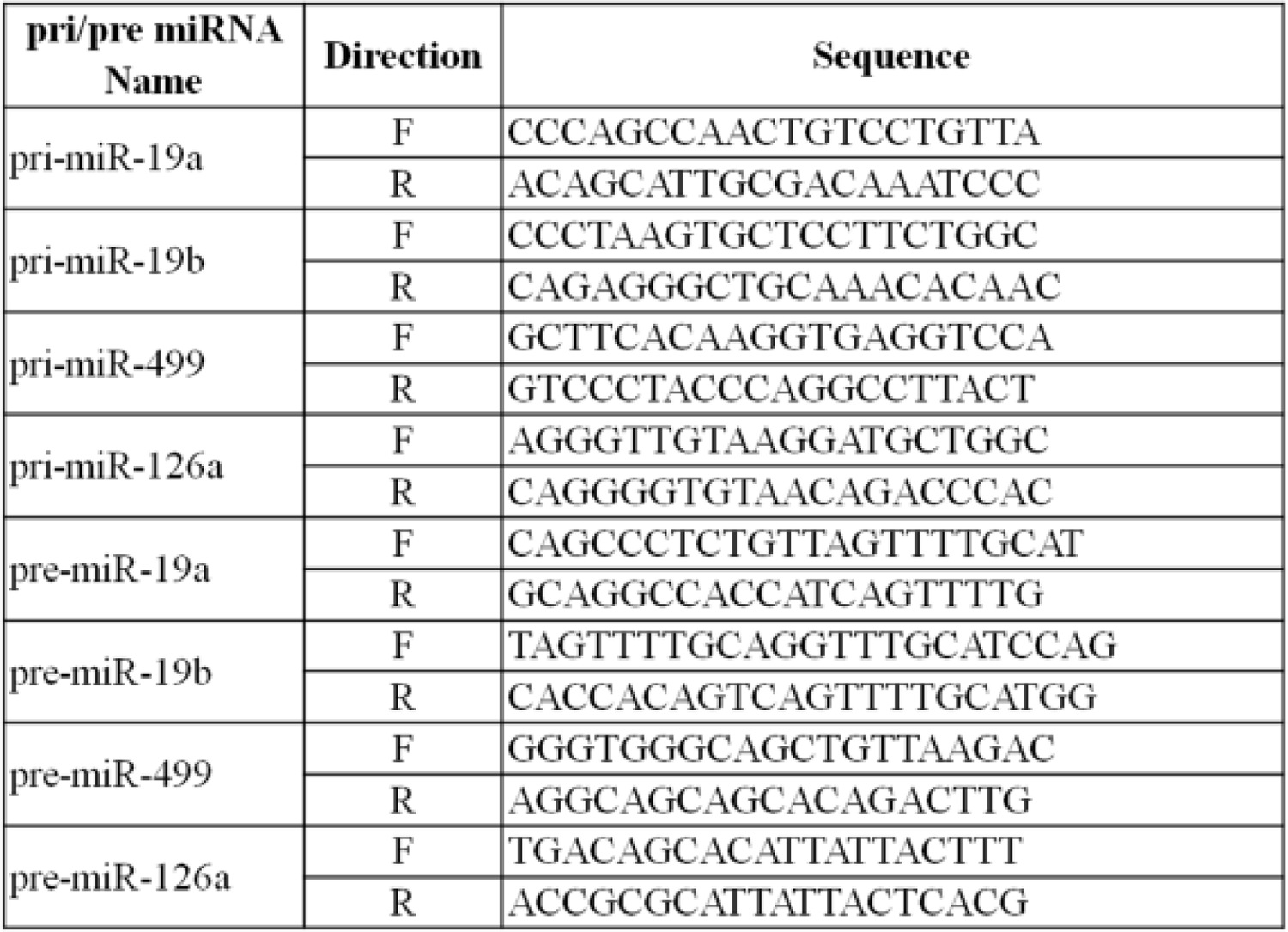
pri and pre-miRNAs’ sequences.

**Supplementary Table 3:**
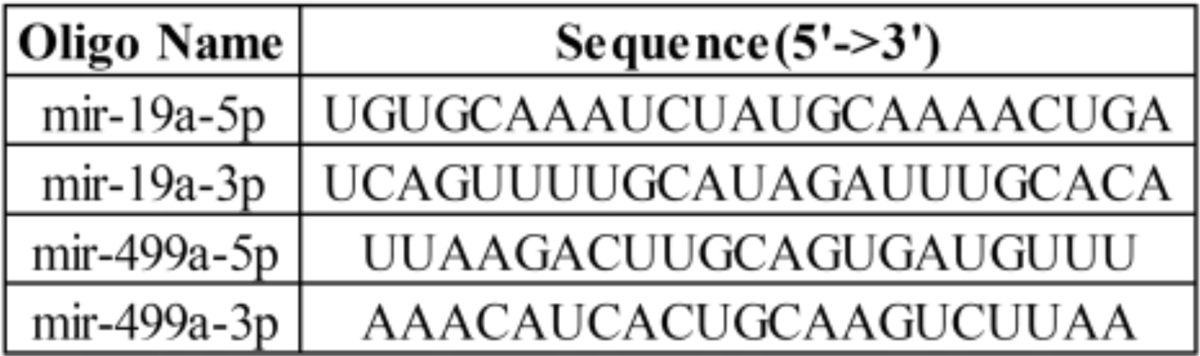
Oligonucleotid’s sequences for microinjection

## METHODS

### Mouse husbandry

Mice were maintained according to the European regulations for the care and use of research animals. The genetic background is *Balb/c*. All animal research was performed in accordance with the relevant guidelines and regulations (Erciyes University animal ethics committee 04-11-2012 12/54).

### RNA microinjection

Oligoribonucleotides synthetic miRNA were adjusted to a concentration of 1 µg/ml and microinjected into *Balb/c* normal fertilized eggs according to established methods of transgenesis^14^. Oligoribonucleotides were obtained from Eurofine (sequences provided in Supplemental Table S3).

Briefly, for obtaining foster mice, healthy female mice and vasectomized male mice were mated. For collecting of embryo, female and male mice were mated at 3 pm. After one night later, mated mice were checked for vaginal plug at 8 am and vaginal plug (+) female mice were selected and divided new cages. Females were sacrificed that afternoon, and their oviducts were enclosed in M2 medium (Sigma, Germany) and embryos were collected under a bino-microscope (Leica, Germany). In order to capture the pronuclei at the most prominent stage, embryos were transferred into M16 medium (Sigma, Germany) with a mouth pipette and incubated at 37°C in 0.5% CO2 (Panasonic, Japan). Then, microinjection of oligonucleotides (1-5 ng/ul) (see in Supplemental Materials Table S3) into the male pronucleus of embryos was performed under an inverted microscope (Nikon, Japan). After the microinjection is finished, the embryos will be transported with a mouth pipette and incubated at 37°C in 0.5% CO2 (Panasonic, Japan). Embryos that died after microinjection under a stereo microscope (Leica, Germany) were separated and surviving embryos were transferred into M2 medium (Sigma, Germany). Living embryos were transferred to foster female mice under anesthesia with a mouth pipette into the oviduct. F0 generation puppies born 21 days after the procedure were fed with breast milk for 21 days, then separated according to their genders and taken into new cages and subjected to behavioral and molecular tests with other groups.

Totally F0 generation 29 mice (5 miR19a-5p microinjected mice, 8 miR-19a-3p microinjected mice, 7 miR-499a-5p microinjected mice and 9 miR-499a-3p), F1 generation 53 mice (19 miR-19a-5p microinjected mice, 15 miR-19a-3p microinjected mice, 12 miR-499a-5p microinjected mice and 10 miR-499a-3p) and 15 healthy control were included to study.

### Behavior Tests

Behavioral experiments were started when miRNA microinjected mice and control group mice were 2 months old. Each mouse underwent a single test daily between 10:00 and 16:00. Only males were tested sequentially on the same day in separate sub-sessions to allow room ventilation and cleanup. The testing tools were cleaned between trials with 70% ethanol and aerated before use. Experiments were videotaped and analyzed offline. Sociability, social-preference and object-recognition and tail suspension tests were analyzed using “EthoVision 9” software (Noldus, Wageningen, Netherlands). Marble-burying tests were analyzed manually by an observer blind to the group of the mice.

#### Novel Object Recognition Test

The device is a square box with an open top, separated by lines at the base and surrounded by a wall. Two identical objects are placed, no larger than the mouse. On the first day, the time the mouse stayed near the objects and the number of visits were calculated. On the second day, one of the objects was changed and a new object was placed. The time the mouse stayed near new and familiar objects, the number of times they went near them, and the discrimination index were measured. When the difference between the total time spent with the new object and the total time spent with the familiar object is divided by the total time and multiplied by 100, the data obtained is called the discrimination index. In summary, what percentage of the total time the mice spent with the new object is when this is measured. This experiment measures the learning and memory functions of mice. In this experiment, the expected behavior of the mice was to spend more time with the new object in order to discover a new object outside of their accustomedness due to their instinct for novelty ^27^

#### Social İnteraction Test

The Social Test assesses cognition in the form of general sociability and interest in social novelty in rodent models of Central Nervous System (CNS) disorders. Rodents normally prefer to spend more time with another rodent (sociality) and will seek out a new stranger more than an acquaintance (social novelty). Crawley’s test of sociability and social novelty consists of a rectangular, three-compartment box. Each room is 19 x 45 cm and is a system with divided walls, a clear plexiglass central chamber that provides free access to each room ^28^. One of the mice is placed in a compartment. While the other compartment is left empty, the mouse originally taken into the experiment was left in the middle compartment for 5 min. Data were created with the EthoVision system in line with the parameters determined to be related to the mouse in the chamber, and statistical analysis was made and compared with the control group.

#### Marble Buried Test

The Marble Buried test is commonly used to measure neophobia (rodent shyness towards new objects, anxiety, obsessive-compulsive or repetitive behaviors) in rodents. The bedding that we normally use in the care of our mice were placed in an empty cage at a height of 5 cm. 20 marbles are arranged in 5 rows with 4 marbles in each row. The experimental mouse was released from one corner of the cage and the mouse was allowed to explore for 30 min. At the end of 30 minutes, the mouse was taken from the cage and the total number of balls buried under the bedding during this period was counted and noted. The number of embedded balls was used for statistical analysis.

#### Tail Suspension Test

The tail hang experiment model of autism is based on observing that after initial escape-directed movements in mice, when placed in an unavoidably stressful situation, they develop a sedentary stance. The stressful situation during tail hanging includes the hemodynamic stress of hanging, and autism models seem to have reduced mobility and escape abilities^29^. The experimental setup was created in such a way that 3 different mice could be tested simultaneously. Thick cardboard-like sheets of 25 cm high were cut and placed between the mice so that the mice could not see each other. Since the mice are white in color, a black background is used in the background. 12 cm long tapes were cut and hung on the experimental setup by sticking them to the tail end of the mouse in such a way that the tails of the mice would not be damaged. Mice were recorded with a video camera for 6 min. By watching the recordings, mobility and immobility time of mice were calculated for 6 min. The immobility time was used for statistical analysis.

After the behavioral experiments of the F0 generation were completed, 2 females and 1 male were mated to obtain the F1 generation. When obtaining the F1 generation, the F0 generation and the control group were sacrificed and blood, hippocampus and sperm samples were taken. For the F1 generation, behavioral experiments were conducted when they were 2 months old. The F1 generation was only studied for behavioral changes and its molecular analyzes were not performed.

### RNA İsolation From Tissue

Hippocampus and blood samples were taken into 500 µl Purezol (Biorad, USA, CA, Cat No: 7326890) and the hippocampus homogenized with a 2 ml syringe. Subsequently, total RNA isolation was carried out according to the manufacturer’s instructions. 200 µl of chloroform was added to the homogenized tissues and blood and centrifuged at +4°C at 12.000 g for 20 min. Then, the aqueous phase was taken and isopropanol was added in a ratio of 1:1. It was centrifuged for 10 min at 12.000 g at +4. The supernatant was poured and 1 ml of 75% ETOH was added then, centrifuged at 7500 g at +4°C. This process was repeated twice, the supernatant was poured and the pellet was incubated at room temperature to dry. 30 µl of nuclease-free water (Qiagen, Germany, Center Mainz, Cat No: 129114) was added and the pellet was resuspended. Total RNAs were stored at −80°C until they started to work.

For sperm RNA isolation, the sperm extracted from the canal taken from the mice were thoroughly turned upside down and shaken in the shaker for 5 minutes and then centrifuged at 1000 rpm for 2 minutes. The supernatant was taken into a new falcon and centrifuged at 3000 rpm for 15 minutes and the supernatant was discarded. 500 µl of dH2O was added, the pellet was resuspended, and after 5 minutes, PBS was added to make up to 5 ml. After centrifugation at 3000 rpm for 15 minutes, the supernatant was discarded. 3 µl DTT and 1000 µl Purezol (Biorad, USA, CA, Cat No: 7326890) were added to each falcon tube and kept on ice for 5 minutes. It was transferred to 1.5 Eppendorf tubes, homogenized with an injector, and 200 µl of chloroform was added. Centrifuging at 12.000 g for 15 minutes at +4, the aqueous phase was taken and isopropanol was added as much as the aqueous phase. It was incubated at −20 for 30 minutes and centrifuged at 12.000 rpm for 30 minutes. The supernatant was discarded, 1 ml of 70% ETOH was added and centrifuged at 7500 rpm for 10 minutes and the supernatant was discarded. 1 ml of 70% ETOH was added and centrifuged at 7500 rpm for 10 minutes. After discarding the supernatant, 500µl dH2O, 25µl 5M NaCl and 1 ml 100% ETOH were added to each sample. It was incubated at −20°C overnight and centrifuged at 10000 rpm for 30 minutes after incubation. The supernatant was poured out. 1 ml of 70% ETOH was added. It was centrifuged at 10000 rpm for 10 minutes. After the supernatant was poured and the remaining alcohol residues were removed, 50 µl of nuclease free water was added and the pellet was resuspended.

### Blood Collection and Serum Separation

Blood samples were collected from the founder mice. Blood samples were collected between 11.00 am and 13.00 to eliminate unwanted parameters. All protocols for serum separation were completed within 1 hour of drawing blood. Serum was separated with centrifugation at 3500 rpm for 10 minutes at room temperature. Hemolyzed samples were excluded from the study. The clear supernatant was collected into new RNase/DNase-free microfuge tubes in 200 μl aliquots. RNA was isolated using High Pure miRNA Isolation Kit (Roche, Mannheim, Germany) according to the manufacturer’s instructions and stored at −80°C until use. All parents gave written informed consent before participation (09-20-2011 committee number: 2011/10).

### DNA-Bound R-loop Hybrid Isolation

DNA-bound R-loop Hybrid were isolated with Quick-DNA/RNA™ Microprep Plus Kit from blood, hippocampus and sperm samples. The manufacturer instructions’ was followed for isolation and DNAse I treatment were applied for taking free-RNA hybrids.

### cDNA Preparation

Isolated RNA samples were reverse-transcribed into cDNA in 20 μl final reaction volumes using miScript II RT Kit(Qiagen,,Germany) as specified in the manufacturer’s protocol. Reverse transcription was performed using the SensoQuest GmbH Thermal Cycler (Göttingen, Germany). cDNA samples were kept at −80°C until PCR analysis.

### Quantitative Real-Time Polymerase Chain Reaction (qRT-PCR)

QRT-PCR was performed by using miScript SYBR® Green PCR Kit (Qiagen, Hilden,Germany Cat No: 218073) with the high-throughput Light Cycler 480 II Real-Time PCR system(Roche, Germany, Mannheim). cDNA samples were diluted with Nuclease Free Water (1:5). The reaction was performed according to manufacturer’s instructions. About 10 μl Syber Green Master Mix, 2 ul 10x Universal Primer, 2 ul primer assays(see in Table S1 and S2) and 4 ul nuclease free water mixed and pipetted into a 96 well plate as 18 ul and 2 μl of 1:5 diluted cDNA pipetted into each well and mixed.. Real-Time PCR step was performed by using Light Cycler 480 II Real-Time PCR system by using the following protocol; thermal mix protocol is followed by Activation step: 95°C for 15 min, denaturation step: 94°C for 15 sec, annealing step: 55°C for 30 sec and extension step: 70°C for 30 sec. After activation step, all steps was carried out for 40 cycles.

### Statistical Analysis

After the results were obtained, comparisons between experimental groups and control groups were made. The compliance of the data to normal distribution was evaluated by the histogram, q-q graphs and Shapiro-Wilk test. Statistical analysis was performed by two-tailed, one-way analysis of variance (ANOVA), uncorrected Fisher’s least significant difference (LSD). Kruskal Walls, student t-test and Mann Whitney-U test depending on whether the data showed normal distribution or not. Data were analyzed using SPSS version 22 (IBM, USA) and Graph-Pad Prism 8.0 software. Results with p values <0.05 were considered statistically significant. Data are expressed as the mean with SD.

